# Predicting Drug Response with Multi-Task Gradient-Boosted Trees in Epilepsy

**DOI:** 10.1101/2025.08.12.669672

**Authors:** Julia Hellmig, Christian M. Boßelmann, Roland Krause, Patrick May, Stefan Wolking, Gianpiero L. Cavalleri, Norman Delanty, John J. Craig, Chantal Depondt, Bobby P.C. Koeleman, Anthony G. Marson, Terence J. O’Brien, Josemir W. Sander, Graeme J. Sills, Pasquale Striano, Federico Zara, Hreinn Stefansson, Kari Stefansson, Sanjay M. Sisodiya, EpiPGX Consortium, Holger Lerche, Nico Pfeifer

## Abstract

**Motivation:** Despite the availability of numerous anti-seizure medications (ASMs), drug resistance remains a major issue for people with epilepsy. The probability of achieving seizure freedom diminishes with each unsuccessful drug trial, and the impact of genetic and clinical markers on ASM response remains unclear. To address this issue, we used state-of-the-art machine learning (ML) methods to predict the response of people with epilepsy to individual ASMs based on their clinical and genomic information.

**Results:** To overcome data sparsity for less common drugs, we implement a multi-task (MT) learning approach for gradient-boosted trees (GBTs), assuming that predicting responses to different ASMs involves similar tasks. This strategy allows models for less prevalent drugs to leverage the more abundant data available for other drugs during training. The proposed model outperforms individual and combined drug-response predictions for most drugs. Our findings identify key genomic and clinical features influencing drug response, enhancing understanding of individual drug responses in people with epilepsy, and aiding clinicians in making informed treatment decisions.

**Availability and Implementation:** Due to privacy reasons data is not publically available. The code will be made available upon acceptance under https://github.com/pfeiferAI/MT-GBT

## 1. Introduction

Epilepsy affects approximately 1% of the global population [9], yet one-third of all people with epilepsy remain drug-resistant despite the availability of over 25 ASMs. Due to reasons that remain unclear, the likelihood of achieving seizure freedom decreases significantly with each treatment attempt [16]. Currently, the treatment choice is based on individual diagnosis, seizure types, other demographic features such as age and sex and cost [31]. With next-generation sequencing (NGS) becoming increasingly affordable, efficient, and widely accessible for clinical use, incorporating an individual’s genetic information into treatment decisions appears to be a promising direction. For the group of genetic-generalized epilepsies (GGE), genetic variants have been shown to play a significant role in disease development [14]. In a genome-wide association study (GWAS), 19 genome-wide significant loci were found to be specific for GGEs [10]. It is assumed that GGEs also have genetic elements that might play a role in drug response [21]. The EpiPGX project, a European FP7-funded consortium, has aimed to address the pharmacogenomics of ASM treatment. Between 2011 and 2015, extensive clinical and genetic data of 12.000 people with epilepsy were collected [31]. Research conducted using this dataset has explored the role of both common and rare genetic variants in drug response. Previous studies relying on traditional GWAS methods have been constrained by limited sample size, resulting in a lack of statistically significant findings. Despite these limitations, initial findings suggest that genetic information may play a crucial role in predicting drug response [18, 30–32]. Few approaches used ML to predict drug response with clinical features or EEG parameters [27, 33]. Only recently supervised ML models were applied to predict the drug-response to brivaracetam based on a combination of genetic and clinical data [5]. This study was limited to the response prediction of one specific ASM, and the limited sample size required rigorous filtering of SNPs prior to model training which may have introduced additional bias. Given that different drugs have different modes of action it is reasonable to assume that clinical and genetic features influencing drug response vary between drugs [13]. This previous approach of individual drug-response prediction was limited by sample size, especially for drugs that are less frequently administered. To efficiently utilize limited data for some drugs, improve performance, and leverage the similarity between certain drugs, more advanced methods are needed. Multi-task learning has been introduced by Caruna et al. stating that tasks that are similar to each other benefit from learning joint representations for prediction [3]. ML methods that frequently have been extended to MT learning frameworks are support vector machines, gaussian processes and (graph) neural networks [2, 3, 6, 11, 25]. Biologically inspired prediction problems that have been approached with MT learning include MHC-I binding prediction, prediction of the functional effects of voltage-gated potassium channel missense variants, splice site prediction and HIV drug suspectibility prediction.[1, 2, 6, 11, 24]. GBTs often show superior prediction performance for tabular datasets when compared to other supervised ML methods [7]. An MT adaptation for GBTs, where feature spaces are shared across tasks has been implemented in Han et al. where they show on simulated datsets that their MT model outperforms the single task model [8]. To our knowledge the combination of MT learning with GBTs has not been applied to any biologically inspired prediction task including drug-response prediction. In this study, we develop ML models that predict the responses of people with epilepsy to individual drugs using both clinical and genetic data. Our findings identify key features influencing drug response and can aid clinicians in making more informed treatment decisions in the future.

## 2. Material and Methods

### 2.1. Clinical Data Preprocessing

Drug-response data was retrieved from the EpiPGX consortium [31]. Participant consent does not allow these data to be made publicly available. Participants were filtered by diagnosis, response, and existing genetic data. Only those with GGE, a clear response or nonresponse label, and existing genotyping data were included in this study. Response is defined as seizure freedom for at least 12 months after start of treatment. Non-response is defined as the recurrence of seizures at 50 percent or more of the frequency before treatment was initiated [31]. We selected treatment trials with the five most prevalent drugs in the dataset, valproate (VPA), lamotrigine (LTG), levetiracetam (LEV), topiramate (TPM), and carbamazepine (CBZ). After filtering the more than 12,000 patients from the EpiPGX dataset, the sample size of treatment trials was reduced to n=416 from 259 individuals. Since an individual can receive several drugs over a course of time, the number of treatment trials is larger than the number of individual patients in the dataset. The clinical features to train the ML models (e.g. age, sex, diagnosis, seizure frequencies and types, age of onset, and the number of ASMs previously tried) were selected with the domain expertise of epileptologists. For every treatment episode, we masked all features revealing information about the ‘future’ e.g. seizure frequency after the treatment episode. A full list of the clinical features used in the models can be found in the Supplement Section B. The continuous clinical features were normalized and centered via Z-score normalization.

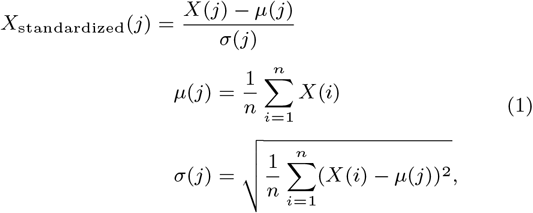

If features were not already encoded appropriately, categorical features were one-hot encoded. Missing values were imputed with the mean of the respective feature. The definition of the label, response or nonresponse, was previously reported in Wolking et al. [31]. Drug-specific datasets were created by filtering the full dataset by the specific drug. For the full dataset, the administered drugs were one-hot encoded and used as features. The numbers of samples in the datasets, the number of responders and nonresponders as well as the gender distribution are summarized in Table 1.

**Table 1.** Summary Statistics of the Data.

| Drug Dataset | # Samples | # Responder | # Non-Responder | Female | Male |
| --- | --- | --- | --- | --- | --- |
| Full | 416 | 184 | 232 | 294 | 122 |
| VPA | 159 | 105 | 54 | 97 | 62 |
| LEV | 83 | 31 | 52 | 60 | 23 |
| LTG | 108 | 29 | 79 | 85 | 23 |
| CBZ | 20 | 6 | 14 | 14 | 6 |
| TPM | 46 | 13 | 33 | 38 | 8 |

### 2.2. Genetic Data Processing

Common SNPs were genotyped by Illumina OmniExpress-12 v1.1 arrays. Preprocessing of the SNPs is described in more detail in Wolking et al. [31]. The gene position of the SNPs with a minor allele frequency *>* 0.01 was annotated with ANNOVAR [28]. For the raw SNP dataset, 626,086 SNPs were filtered based on their gene annotation. The lists of genes and gene sets were derived from previous publications [14, 31, 32] and contain genes related to epilepsy, ion channels, the GABAergic and glutamatergic pathways and other neuronal pathways, as well as genes responsible for absorption, distribution, metabolization, and, excretion of drugs (ADME genes) and can be found in Section C in the Supplement. By only keeping SNPs that are located in the genes that are part of the gene sets (see Supplement Section C), the number of SNPs was reduced to 52,530. For the raw-SNPs dataset, filtered SNPs were encoded with 0, 1 and 2 for homozygote major allele, heterozygote and homozygote minor allele, respectively. The gene counts dataset consists of the counts of variants in each gene for each individual. Duplicate genes from the gene sets were removed. For this, we added up the SNP encodings per gene. Genes can belong to more than one gene set. To create the gene set dataset, the counts of variants per gene set were calculated. The gene set and gene counts dataset were normalized using Z-score normalization (see equation 1). The numbers of features for each dataset are summarized in Table 2.

**Table 2.** Number of features. As the drug name of the treatment trial is one hot encoded in the dataset for all drugs, this specific dataset contains five more clinical features than the drug-specific datasets (shown in brackets).

| Dataset | No. Features | Clinical | Genomic |
| --- | --- | --- | --- |
| Clinical | 44 (49) | 44 (49) | 0 |
| Raw SNPs | 52,574 (52,570) | 44 (49) | 52,530 |
| Gene counts | 1,811 (1816) | 44 (49) | 1,767 |
| Gene sets | 69 (74) | 44 (49) | 25 |

### 2.3. Model Training

We trained four different supervised ML algorithms to predict drug response for ASMs: a logistic regression model, a random forest classifier, SMVs with different kernel functions (polynomial kernel of degree two and three, a linear kernel, and a radial basis function kernel) and GBTs.

For all models, we applied a nested, stratified 5×5-fold cross-validation for hyperparameter fitting (inner folds) and performance evaluation (outer folds). As we are dealing with imbalanced datasets, the performance of the models was evaluated on the test sets with the area under the precision-recall curve value (AUPRC) and its standard deviation (SD).

### 2.4. Feature Importance

Feature importance was calculated from the XGBoost trees with the built-in feature importance method. The importance is calculated by the average gain of all splits in which the feature is used. Features were sorted by their importance in descending order to report the top ten most important features per dataset. The directions of feature importance were calculated for categorical and continuous features.

### 2.5. MT-GBT

We adapted the MT approach for GBTs from Han et al. [8]. The idea of this approach is to learn a separate model of GBTs for each task but include information from other tasks by generating global and task-specific feature sets. The influence of task-specific (Ω_*t*_) and the global feature set (Ω_*G*_) can be controlled with two penalty parameters *μ*_*t*_ and *μ*_*g*_, respectively. The loss function that is minimized for feature j and threshold *s*_*j*_ is:

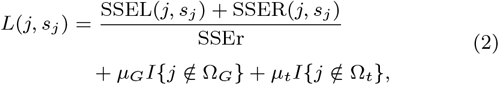

with

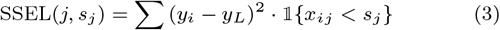

and

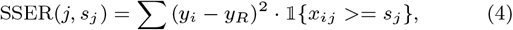

as the sum of squared errors for the left and right child tree, where ⊮ is the indicator function for the case that the value of feature j of sample *x*_*i*_ is smaller (SSEL) or larger than/ equal to *s*_*j*_ (SSER) and *y*_*R*_, *y*_*L*_ being the mean values of the right and left node, respectively [8]. The additional term in the loss function causes features that are not in the task-specific or group feature set to be penalized with the respective penalty. Therefore, introducing new features as splits becomes more costly and the reusing of features already contained in the task-specific feature set and/or group feature set is preferred. In our setting different tasks are predicting drug response for the five individual ASMs. The MT-GBT algorithm is described in Algorithm 1. We trained the model on the clinical and gene sets dataset and for different *μ*_*t*_ and *μ*_*g*_ combinations. Performance was reported with the AUPRC score and its SD over a 5×5-fold nested, stratified cross-validation.

#### Algorithm 1

Pseudocode for MT gradient-boosted trees

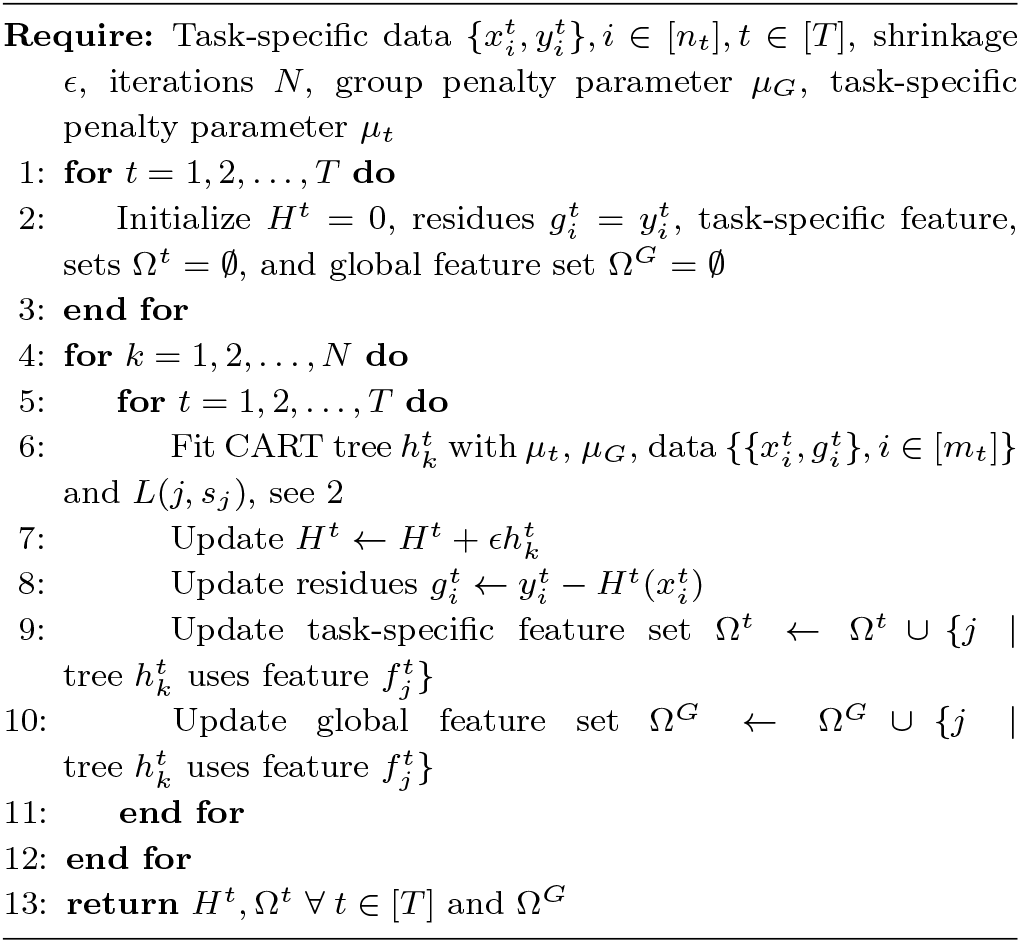

### 2.6. Implementation

The logistic regression, random forest, and SVM models were implemented with the scikit learn package in Python 3.8. For the logistic regression model we used the default lbfgs solver with the l1 penalty. The measure of impurity used for the random forest classifier was the gini-impurity. [22]. For the single-task GBTs model we used the XGBoost package in Python 3.8 with the binary logistic learning-objective. [4]. We implemented the MT GBT model by extending the Classification and Regression Trees (CART) algorithm with a custom loss function described in more detail in Section 2.5. The code will be made available upon acceptance under https://github.com/pfeiferAI/MT-GBT.

## 3. Results

### 3.1. Supervised ML models trained on clinical data

First, we used the clinical dataset for all drugs to predict drug response using four different supervised ML models. We report the performance of the model with the AUPRC in Table 3. Among these models, GBTs and random foreests significantly outperformed the other models. We continued all following experiments with GBTs.

**Table 3.** AUPRC performances of four supervised learning models on the clinical dataset for all drugs. Four different ML models were trained on the clinical dataset for all drugs.

| Model | AUPRC |
| --- | --- |
| SVM (Poly 3 kernel) | $0.56 \pm 0.00$ |
| GBTs | **$0.63 \pm 0.04$** |
| Random Forests | **$0.64 \pm 0.06$** |
| Logistic Regression | $0.43 \pm 0.03$ |

### 3.2. Prediction of drug response for individual drugs

We assessed how the prediction performance of the model changes when predicting the response to individual ASMs. We split the dataset by drugs, resulting in drug-specific datasets for the five most prevalent drugs in the dataset. GBTs were trained on the drug-specific datasets in the same way as on the complete dataset (see 2.3). We found that drug-specific response prediction for VPA improved substantially when compared to the full clinical dataset (Table 4). Meanwhile, the models for CBZ and TPM performed poorly likely due to small sample sizes.

**Table 4.** AUPRC performances with SD for GBTs. GBTs were trained on all drugs and drug-specific datasets with different types of genetic information added to the dataset. The AUPRC performances are reported with their SD on the test sets of a stratified 5-fold nested cross-validation.

| Metric | All | VPA | LEV | LTG | CBZ | TPM |
| --- | --- | --- | --- | --- | --- | --- |
| Random Baseline | 0.56 | 0.34 | 0.63 | 0.73 | 0.72 | 0.7 |
| Clinical | $0.63 \pm 0.04$ | $0.43 \pm 0.1$ | $0.61 \pm 0.03$ | **$0.79 \pm 0.04$** | $0.72 \pm 0.16$ | $0.72 \pm 0.07$ |
| Clinical + Raw SNPs | $0.59 \pm 0.05$ | $0.3 \pm 0.02$ | **$0.67 \pm 0.11$** | $0.67 \pm 0.02$ | $0.68 \pm 0.07$ | **$0.75 \pm 0.04$** |
| Clinical + Gene counts | $0.62 \pm 0.02$ | **$0.46 \pm 0.06$** | $0.62 \pm 0.05$ | $0.72 \pm 0.02$ | **$0.83 \pm 0.19$** | **$0.75 \pm 0.07$** |
| Clinical + Gene sets | **$0.64 \pm 0.04$** | $0.4 \pm 0.05$ | $0.64 \pm 0.06$ | $0.74 \pm 0.02$ | $0.69 \pm 0.10$ | $0.69 \pm 0.05$ |

### 3.3. Improving drug-response prediction with genetic features

Next, we investigated whether the addition of genetic information improves the predictive performance of our models. Therefore, we trained GBTs on three datasets that combined genetic and clinical information. Genetic information was added in three different forms, genotyping information, counts of variants per gene (gene counts) and counts of variants per gene set (gene set counts). The AUPRC and Matthews correlation coefficient (MCC) scores with their SDs are reported in Table 4 and Supplement Section E, respectively. For all drugs except for LTG the addition of genetic data in some form improved the prediction performance of the model. When predicting response across all drugs the accumulation of genetic information to gene set counts improved the performance of the model compared to prediction with raw SNPs (AUPRC 0.64±0.04 vs. 0.59±0.05). There is no clear picture which representation of genetic information is most beneficial for drug-response prediction. Interestingly, drug-response prediction for LTG did not benefit from the addition of genetic information, suggesting that clinical data was sufficient. The largest performance improvements when predicting individual drug response—compared to joint modeling across all drugs—are observed for VPA and CBZ, particularly when using gene counts as features. This suggests that these drugs may have more distinct, drug-specific genetic predictive signals, especially in the case of CBZ, which are better captured when models are trained separately rather than jointly.

### 3.4. Feature Importance of GBTs

Thus, we observed that the impact of genetic and clinical features on prediction performance varied across drugs. To investigate biological plausibility and possible underlying mechanisms, we evaluated the ten most important features of the gradient-boosted tree models. The validity of feature importance estimates depends strongly on model performance. Thus, we report only the two most robust models: those for all drugs and VPA, respectively in Fig. 1. It is important to note that these feature importance estimates do not establish causal relationships between the features and model predictions. The clinical features that contributed most to the models were 1) the age at the first seizure, 2) the number of treatment trials, 3) seizure types, and 4) a family history of epilepsy. These findings align well with clinical experience and previous literature, where each feature is known to directly impact the likelihood of seizure freedom [17, 23]. Further, gene sets encoding for components of GABAergic transmission were among the most important features for all drugs. This may be on the one hand due to the reported modification of of GABAergic signalling by VPA, TPM, and to a lesser extent, LTG [12, 15, 20, 26, 29] and on the other hand to the specific involvement of GABAergic genes in the pathogenesis of GGEs [14]. The latter would be in line with our observation that gene sets containing genes of known relevance in generalized epilepsies show to have great influence in VPA response predicition. Why this affects mainly the VPA response may have to do with the superior response of VPA compared to other drugs in GGE in general [12].

**Fig. 1:**
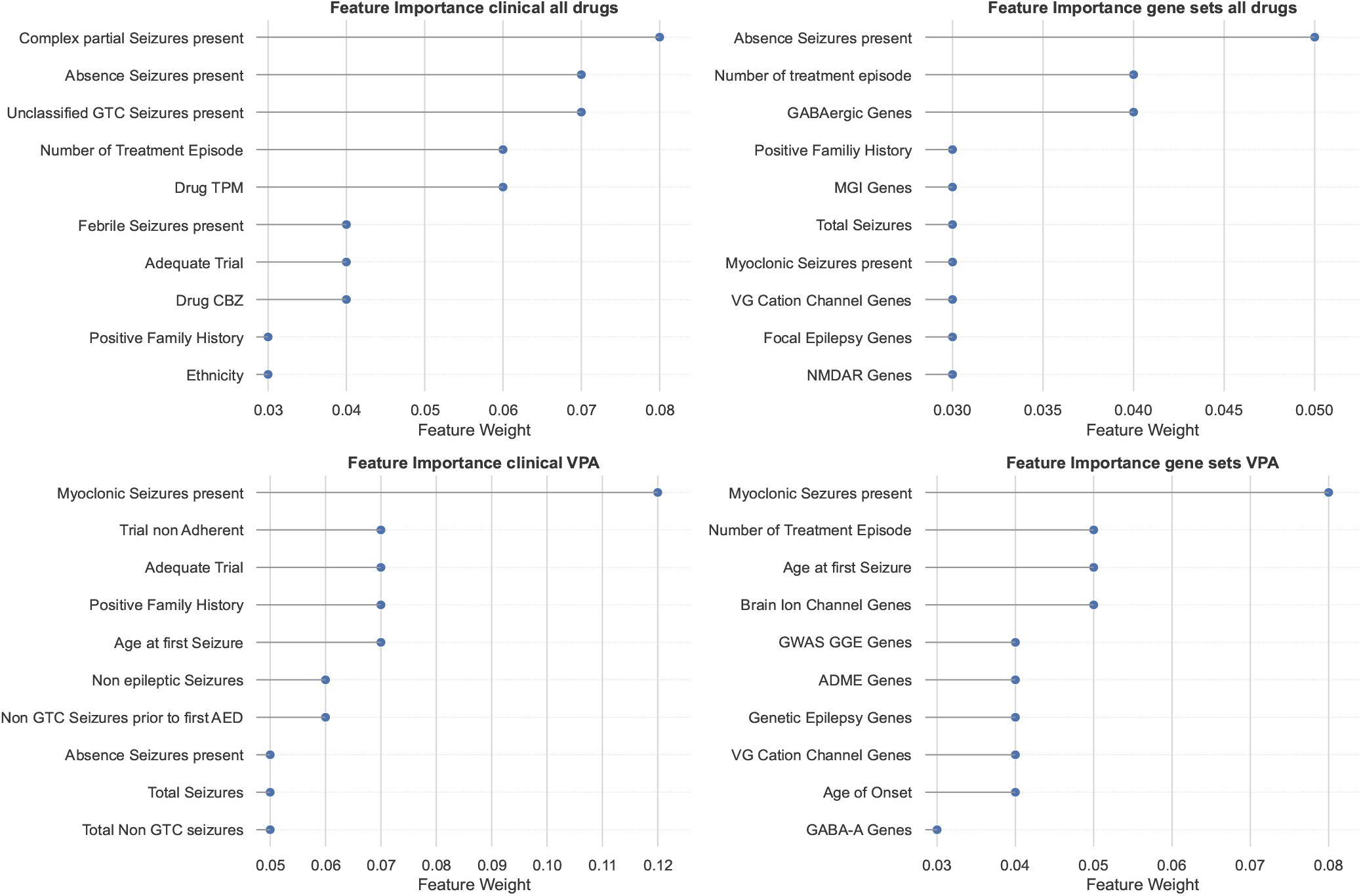
Feature Importance of GBTs. The feature importance of the top 10 features for the clinical dataset for A) all drugs, B) VPA and the gene set dataset for C) all drugs and D) VPA is shown. The features are sorted in descending order by their absolute importance. Importance was calculated with the built-in feature performance function of XGBoost.

### 3.5. MT learning improves prediction performance for most drugs

To this point we have shown, that the utility of clinical or genetic features, varies considerably across drugs given their different underlying mechanisms. We hypothesized that MT learning may enable more accurate predictions by accounting for these differences, resulting in efficient prediction across all drugs. We applied our MT-GBT approach to three datasets, namely the clinical, the gene counts and the gene sets datasets. Given our computational resources it was not possible to apply the model to the raw SNPs dataset. No parameter settings of *μ*_*t*_ and *μ*_*g*_ were optimal across all drugs. This indicates that the optimal amount of data sharing across tasks differs between individual drugs. Model performances of the stratified nested cross-validation are shown in Fig. 2. MT learning improved the prediction performance for all drugs except LTG. Notably, for TPM, which has a small sample size, performance increased significantly due to shared learning across tasks. With our method it is possible to get further insights into task-specific and global feature sets. The intersections of those feature sets are plotted with upset plots and can be found in Supplement Section F [19]. Clinical features are mainly shared between all tasks, however, with increasing amounts of genetic information used for training, features became more drug-specific.

**Fig. 2:**
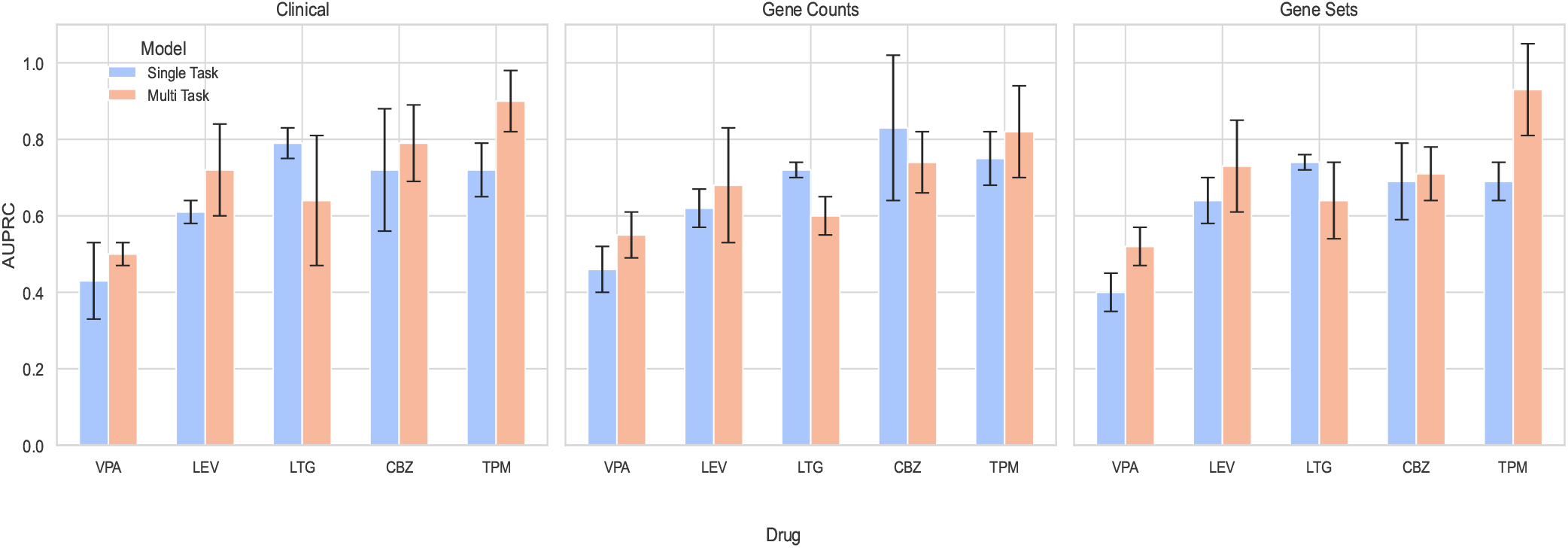
Performance comparison of MT GBTs and single-task learning across three datasets. The Figure shows the area under the precision recall curve of MT GBTs and single-task learning across three datasets with their SD: clinical dataset, gene counts, and gene sets. For each dataset, five individual drug models were trained. The single-task models were trained with XGBoost gradient-boosted decision trees. Those findings were already reported in Table 3. For the drugs VPA, LEV and TPM the MT approach outperformed the single task learning approach in all datasets. LTG did not benefit from MT learning.

## 4. Expected clinical impact

The ability to predict individual drug response prior to treatment initiation could significantly enhance clinical decision-making by reducing time spent on ineffective medications, minimizing side effects, and accelerating seizure control. By integrating readily available clinical and genetic data, the proposed model offers a pragmatic tool that could be embedded into routine workflows. Importantly, the prediction pipeline is computationally efficient, allowing for rapid output generation, which supports its feasibility in real-world settings. Such a model has the potential to improve cost-effectiveness by optimizing treatment earlier in the care pathway and reducing the need for prolonged trial-and-error medication strategies.

## 5. Potential bias and limitations

Several factors temper the generalizability of our findings. First, the retrospective design introduces inherent selection bias and limits control over data completeness. Second, we did not have systematic measures of medication adherence; non-adherence could confound response labels and inflate misclassification rates. Third, follow-up duration varied widely (6–36 months), potentially masking late responders or non-responders and adding noise to the binary outcome. Finally, although our sample size (n = 416 treatment episodes) is larger than in prior ML studies, it remains modest for high-dimensional genomic analyses, and some drugs (e.g., CBZ) still have limited representation, which may affect feature stability. Prospective, multicentre validation with uniform follow-up and adherence monitoring will be essential to confirm clinical utility.

## 6. Conclusion

In this study, we explored the applicability of ML methods for predicting drug responses in individuals with epilepsy. Among various supervised learning models, GBTs emerged as the most effective. To further assess model performance, we incorporated genetic data in various formats alongside clinical features. Incorporating genetic information leads to improved performance. Additionally, the feature-importance evaluation reveals that genetic features consistently rank among the top ten most important factors in most models. This indicates that incorporating genetic information is valuable for predicting drug response. However, it is crucial to strike a balance. On the one hand, having too many features, such as raw SNPs, can overwhelm the model without adding meaningful value. On the other hand, excessive aggregation, like gene set counts, can dilute the quality of the information. Including both raw genetic variants and more sophisticatedly aggregated features, such as pathway- or disease-specific polygenic risk scores, could add depth to the genetic data used in ML models. Filtering the features to retain only those already known to be relevant for epilepsy helps reduce noise and eliminates irrelevant data. Finding the right trade-off between feature quantity, aggregation, and filtering is key to optimizing model performance. Additionally, employing unsupervised feature selection methods or loosening pre-filtering constraints could uncover novel genetic features not yet identified in the context of epilepsy. Training models on drug-specific datasets shows that for some drugs it is beneficial to learn drug response solely on data from their own drug. This is reasonable, as the modes of action between drugs differ significantly. However, for drugs that are not that frequently administered, the sample size is so small that learning an ML model only on drug-specific data is not feasible. We adapted an MT learning framework for GBTs to address this issue. Our results show that sharing information across tasks is beneficial for nearly all drugs, with the exception of LTG, which benefits the most from being learned independently in a specific dataset. This approach could be further refined by adjusting the extent of knowledge sharing between drugs, depending on their similarity.

## 8. Author contributions statement

J.H., N.P., S.W., R.K., P.M. and H.L. conceived the experiments, R.K., P.M., S.W., G.C., N.D., J.C., C.D., B.K., A.M., T.B., J.S., G.S., P.S., F.Z., H.S., K.S., S.S. and H.L. curated the data, J.H. conducted the experiments, J.H., C.B., N.P. and H.L. analysed the results. J.H. and C.B. wrote and reviewed the manuscript. All authors revised the manuscript for scientific content.

## References

1. S. Bickel, J. Bogojeska, T. Lengauer, and T. Scheffer. Multi-task learning for hiv therapy screening. In Proceedings of the 25th international conference on Machine learning, pages 56–63, 2008.

2. C. M. Boßelmann, U. B. Hedrich, P. Müller, L. Sonnenberg, S. Parthasarathy, I. Helbig, H. Lerche, and N. Pfeifer. Predicting the functional effects of voltage-gated potassium channel missense variants with multi-task learning. EBioMedicine, 81, 2022.

3. R. Caruana. Algorithms and applications for multitask learning. In ICML, pages 87–95. Citeseer, 1996.

4. T. Chen and C. Guestrin. Xgboost: A scalable tree boosting system. In Proceedings of the 22nd acm sigkdd international conference on knowledge discovery and data mining, pages 785–794, 2016.

5. J. de Jong, I. Cutcutache, M. Page, S. Elmoufti, C. Dilley, H. Fröhlich, and M. Armstrong. Towards realizing the vision of precision medicine: Ai based prediction of clinical drug response. Brain, 144(6):1738–1750, 2021.

6. M. Gönen and A. A. Margolin. Drug susceptibility prediction against a panel of drugs using kernelized bayesian multitask learning. Bioinformatics, 30(17):i556–i563, 2014.

7. L. Grinsztajn, E. Oyallon, and G. Varoquaux. Why do tree-based models still outperform deep learning on typical tabular data? Advances in neural information processing systems, 35: 507–520, 2022.

8. C. Han, N. Rao, D. Sorokina, and K. Subbian. Scalable Feature Selection for (Multitask) Gradient Boosted Trees. Proceedings of the 23rdInternational Conference on Artificial Intelligence and Statistics (AISTATS), 2020.

9. D. Hesdorffer, G. Logroscino, E. Benn, N. Katri, G. Cascino, and W. Hauser. Estimating risk for developing epilepsy: a population-based study in rochester, minnesota. Neurology, 76(1):23–27, 2011.

10. International League Against Epilepsy Consortium on Complex Epilepsies, R. Stevelink, C. Campbell, and S. e. a. Chen. GWAS meta-analysis of over 29,000 people with epilepsy identifies 26 risk loci and subtype-specific genetic architecture. Nature Genetics, 55(9):1471–1482, Sept. 2023. ISSN 1061-4036, 1546-1718. doi: 10.1038/s41588-023-01485-w.

11. L. Jacob and J.-P. Vert. Efficient peptide–mhc-i binding prediction for alleles with few known binders. Bioinformatics, 24(3):358–366, 2008.

12. P. Klein, R. M. Kaminski, M. Koepp, and W. Löscher. New epilepsy therapies in development. Nature Reviews Drug Discovery, 23(9):682–708, 2024.

13. J. K. Knowles, I. Helbig, C. S. Metcalf, L. Lubbers, L. L. Isom, S. Demarest, and et al. Precision medicine for genetic epilepsy on the horizon: Recent advances, present challenges, and suggestions for continued progress. Epilepsia, 63(10): 2461–2475, Oct. 2022. ISSN 0013-9580, 1528-1167. doi: 10.1111/epi.17332.

14. M. Koko, R. Krause, T. Sander, and D. R. e. a. Bobbili. Distinct gene-set burden patterns underlie common generalized and focal epilepsies. eBioMedicine, 72:103588, Oct. 2021. ISSN 23523964. doi: 10.1016/j.ebiom.2021.103588.

15. R. Kuzniecky, S. Ho, J. Pan, R. Martin, F. Gilliam, E. Faught, and H. Hetherington. Modulation of cerebral gaba by topiramate, lamotrigine, and gabapentin in healthy adults. Neurology, 58(3):368–372, 2002.

16. P. Kwan and M. J. Brodie. Early identification of refractory epilepsy. New England Journal of Medicine, 342(5):314–319, 2000.

17. P. Kwan, S. C. Schachter, and M. J. Brodie. Drug-resistant epilepsy. New England Journal of Medicine, 365(10):919–926, 2011.

18. C. Leu, A. Avbersek, R. Stevelink, H. M. Custodio, S. Chen, D. Speed, and et al. Genome-wide association meta-analyses of drug-resistant epilepsy. EBioMedicine, 115, 2025.

19. A. Lex, N. Gehlenborg, H. Strobelt, R. Vuillemot, and H. Pfister. UpSet: Visualization of Intersecting Sets. IEEE Transactions on Visualization and Computer Graphics, 20 (12):1983–1992, Dec. 2014. ISSN 1077-2626. doi: 10.1109/TVCG.2014.2346248.

20. W. Löscher. Valproate enhances gaba turnover in the substantia nigra. Brain research, 501(1):198–203, 1989.

21. W. Löscher, U. Klotz, F. Zimprich, and D. Schmidt. The clinical impact of pharmacogenetics on the treatment of epilepsy. Epilepsia, 50(1):1–23, 2009.

22. F. Pedregosa, G. Varoquaux, A. Gramfort, V. Michel, B. Thirion, O. Grisel, and et al. Scikit-learn: Machine learning in Python. Journal of Machine Learning Research, 12:2825–2830, 2011.

23. A. Pitkänen, D. C. Henshall, J. H. Cross, R. Guerrini, S. Jozwiak, M. Kokaia, and et al. Advancing research toward faster diagnosis, better treatment, and end of stigma in epilepsy. Epilepsia, 60(7):1281–1292, 2019.

24. G. Schweikert, G. Rätsch, C. Widmer, and B. Schölkopf. An Empirical Analysis of Domain Adaptation Algorithms for Genomic Sequence Analysis.

25. C. Song, X. Liu, Z. Xiong, L. Liu, and W. Zhang. Gnndrp: Graph neural network with multi-task learning for drug response prediction. IEEE Transactions on Computational Biology and Bioinformatics, 22(3):1052–1059, 2025. doi: 10.1109/TCBBIO.2025.3548692.

26. D. M. Treiman. Gabaergic mechanisms in epilepsy. Epilepsia, 42(3):8–12, 2001. doi: 10.1046/j.1528-1157.2001.042suppl.3008.x.

27. B. Wang, X. Han, Z. Zhao, N. Wang, P. Zhao, M. Li, and et al. EEG-Driven Prediction Model of Oxcarbazepine Treatment Outcomes in Patients With Newly-Diagnosed Focal Epilepsy. Frontiers in Medicine, 8:781937, Jan. 2022. ISSN 2296-858X. doi: 10.3389/fmed.2021.781937.

28. K. Wang, M. Li, and H. Hakonarson. Annovar: functional annotation of genetic variants from high-throughput sequencing data. Nucleic Acids Research, 38(16):e164–e164, 07 2010. ISSN 0305-1048. doi: 10.1093/nar/gkq603.

29. R. D. Whitlow, A. Sacher, D. D. Loo, N. Nelson, and S. Eskandari. The anticonvulsant valproate increases the turnover rate of γ-aminobutyric acid transporters. Journal of Biological Chemistry, 278(20):17716–17726, 2003.

30. S. Wolking, C. Moreau, A. T. Nies, E. Schaeffeler, M. McCormack, P. Auce, and et al. Testing association of rare genetic variants with resistance to three common antiseizure medications. Epilepsia, 61(4):657–666, Apr. 2020. ISSN 0013-9580, 1528-1167. doi: 10.1111/epi.16467.

31. S. Wolking, H. Schulz, A. T. Nies, M. McCormack, E. Schaeffeler, P. Auce, and et al. Pharmacoresponse in Genetic Generalized Epilepsy: A Genome-Wide Association Study. Pharmacogenomics, 21(5):325–335, Apr. 2020. ISSN 1462-2416, 1744-8042. doi: 10.2217/pgs-2019-0179.

32. S. Wolking, C. Campbell, C. Stapleton, M. McCormack, N. Delanty, C. Depondt, and et al. Role of Common Genetic Variants for Drug-Resistance to Specific Anti-Seizure Medications. Frontiers in Pharmacology, 12:688386, June 2021. ISSN 1663-9812. doi: 10.3389/fphar.2021.688386.

33. L. Yao, M. Cai, Y. Chen, C. Shen, L. Shi, and Y. Guo. Prediction of antiepileptic drug treatment outcomes of patients with newly diagnosed epilepsy by machine learning. Epilepsy & Behavior, 96:92–97, July 2019. ISSN 15255050. doi: 10.1016/j.yebeh.2019.04.006.

