## Supplementary Information for "Predicting Drug Response with Multi-Task Gradient-Boosted Trees in Epilepsy"

for

### A. EpiPGX Consortium Members:

EpiPGX Consortium Andreja Avbersek, Costin Leu, Kristin Heggeli, Rita Demurtas, Joseph Willis, Doug Speed, Narek Sargsyan, Krishna Chinthapalli, Mojgansadat Borghei, Antonietta Coppola, Antonio Gambardella, Stefan Wolking, Felicitas Becker, Sarah Rau, Christian Hengsbach, Yvonne G. Weber, Bianca Berghuis, Wolfram S. Kunz, Mark McCormack, Norman Delanty, Ellen Campbell, Lárus J. Gudmundsson, Andrés Ingason, Kári Stefánsson, Reinhard Schneider, Rudi Balling, Pauls Auce, Ben Francis, Andrea Jorgensen, Andrew Morris, Sarah R. Langley, Prashant K. Srivastava, Martin J. Brodie, Marian Todaro, Slave Petrovski, Jane L. Hutton, Fritz Zimprich, Martin Krenn, Hiltrud Muhle, Karl M. Klein, Rikke S. Møller, Marina Nikanorova, Sarah Weckhuysen, Zvonka Rener-Primec, Gianpiero L. Cavalleri, John Craig, Chantal Depondt, Michael R. Johnson, Bobby P. C. Koeleman, Roland Krause, Holger Lerche, Anthony G. Marson, Terence J. O'Brien, Josemir W. Sander, Graeme J. Sills, Hreinn Stefansson, Pasquale Striano, Federico Zara, Sanjay M. Sisodiya

### B. Clinical features

List of clinical features used for drug-response prediction.

- Drug trial non-adherent
- Gender
- Ethnicity
- Diagnosis
- Age at diagnosis
- Age at first seizure
- Hippocampal Sclerosis (right, left)
- Family History
- Neurological exam results
- Neurological progressive disorder
- Number of treatment episodes
- Seizure types:
  - Absence
  - Clonic
  - Tonic
  - Myoclonic

- Atonic
  - Febrile
  - Simple Partial
  - Complex Partial
  - Unclassified Partial
  - Unclassified GTC
  - Uncertain epileptic
  - Non-epileptic
  - Secondarily GTC
- Seizure frequencies (Categorical and Absolute values):
  - Number of GTC/Non-GTC, total seizures prior to first AED

### C. Gene Sets

List of gene sets and genes used for drug-response predictions.

| **Gene Set** | **Genes in Set** |
|---|---|
| VG-Cation genes | CACNA1A, CACNA1B, CACNA1C, CACNA1D, CACNA1E, CACNA1F, CACNA1G, CACNA1H, CACNA1I, CACNA1S, CACNA2D1, CACNA2D2, CACNA2D3, CACNA2D4, CACNB1, CACNB2, CACNB3, CACNB4, HCN1, HCN2, HCN3, HCN4, KCNA1, KCNA10, KCNA2, KCNA3, KCNA4, KCNA5, KCNA6, KCNA7, KCNAB1, KCNAB2, KCNAB3, KCNB1, KCNB2, KCNC1, KCNC2, KCNC3, KCNC4, KCND1, KCND2, KCND3, KCNE1, KCNE1L, KCNE2, KCNE3, KCNE4, KCNF1, KCNG1, KCNG2, KCNG3, KCNG4, KCNH1, KCNH2, KCNH3, KCNH4, KCNH5, KCNH6, KCNH7, KCNH8, KCNQ1, KCNQ2, KCNQ3, KCNQ4, KCNQ5, KCNRG, KCNS1, KCNS2, KCNS3, KCNT1, KCNV1, KCNV2, SCN10A, SCN11A, SCN1A, SCN1B, SCN2A, SCN2B, SCN3A, SCN3B, SCN4A, SCN4B, SCN5A, SCN7A, SCN8A, SCN9A. |
| GABA-A genes | GABRA1, GABRA2, GABRA3, GABRA4, GABRA5, GABRA6, GABRB1, GABRB2, GABRB3, GABRD, GABRE, GABRG1, GABRG2, GABRG3, GABRP, GABRQ, GABRR1, GABRR2. |
| GABAergic genes | ABAT, ADCY1, ADCY2, ADCY3, ADCY4, ADCY5, ADCY6, ADCY7, ADCY8, ADCY9, ANK2, ANK3, ARHGEF9, DISC1, DLC1, DNAI1, FGF13, GABARAP, GABARAPL1, GABARAPL2, GABBR1, GABBR2, GABRA1, GABRA2, GABRA3, GABRA4, GABRA5, GABRA6, GABRB1, GABRB2, GABRB3, GABRD, GABRE, GABRG1, GABRG2, GABRG3, GABRP, GABRQ, GABRR1, GABRR2, GAD1, GAD2, GLS, GLS2, GLUL, GNAI1, GNAI2, GNAI3, GNAO1, GNB1, GNB2, GNB3, GNB4, GNB5, GNG10, GNG11, GNG12, GNG13, GNG2, GNG3, GNG4, GNG5, GNG7, GNG8, GNGT1, GNGT2, GPHN, HAP1, KCNB2, KCNC1, KCNC2, KCNC3, KCNJ6, KIF5A, KIF5B, KIF5C, MAGI1, MKLN1, MTOR, MYO5A, NLGN2, NRXN1, NSF, PFN1, PLCL1, PRKACA, PRKACB, PRKACG, PRKCA, PRKCB, PRKCG, RDX, SCN1A, SCN1B, SCN2B, SCN3A, SCN8A, SEMA4D, SLC12A2, SLC12A5, SLC32A1, SLC38A1, SLC38A2, SLC38A5, SLC6A1, SLC6A11, SLC6A13, SRC, STARD13, TRAK1, TRAK2. |

| | |
|---|---|
| Ion channel genes | CACNA1A, CACNA1B, CACNA1C, CACNA1D, CACNA1E, CACNA1F, CACNA1G, CACNA1H, CACNA1I, CACNA1S, CATSPER1, CATSPER2, CATSPER3, CATSPER4, CHRNA1, CHRNA10, CHRNA2, CHRNA3, CHRNA4, CHRNA5, CHRNA6, CHRNA7, CHRNA9, CHRNB1, CHRNB2, CHRNB3, CHRNB4, CHRND, CHRNE, CHRNG, CNGA1, CNGA2, CNGA3, CNGA4, CNGB1, CNGB3, GABRA1, GABRA2, GABRA3, GABRA4, GABRA5, GABRA6, GABRB1, GABRB2, GABRB3, GABRD, GABRE, GABRG1, GABRG2, GABRG3, GABRP, GABRQ, GABRR1, GABRR2, GLRA1, GLRA2, GLRA3, GLRA4, GLRB, GRIA1, GRIA2, GRIA3, GRIA4, GRID1, GRID2, GRIK1, GRIK2, GRIK3, GRIK4, GRIK5, GRIN1, GRIN2A, GRIN2B, GRIN2C, GRIN2D, GRIN3A, GRIN3B, HCN1, HCN2, HCN3, HCN4, HTR3A, HTR3B, HTR3C, HTR3D, HTR3E, KCNA1, KCNA10, KCNA2, KCNA3, KCNA4, KCNA5, KCNA6, KCNA7, KCNB1, KCNB2, KCNC1, KCNC2, KCNC3, KCNC4, KCND1, KCND2, KCND3, KCNF1, KCNG1, KCNG2, KCNG3, KCNG4, KCNH1, KCNH2, KCNH3, KCNH4, KCNH5, KCNH6, KCNH7, KCNH8, KCNJ1, KCNJ10, KCNJ11, KCNJ12, KCNJ13, KCNJ14, KCNJ15, KCNJ16, KCNJ2, KCNJ3, KCNJ4, KCNJ5, KCNJ6, KCNJ8, KCNJ9, KCNK1, KCNK10, KCNK12, KCNK13, KCNK15, KCNK16, KCNK17, KCNK18, KCNK2, KCNK3, KCNK4, KCNK5, KCNK6, KCNK7, KCNK9, KCNMA1, KCNN2, KCNN3, KCNN4, KCNQ1, KCNQ2, KCNQ3, KCNQ4, KCNQ5, KCNS1, KCNS2, KCNS3, KCNT1, KCNT2, KCNU1, KCNV1, KCNV2, MCOLN1, MCOLN2, MCOLN3, P2RX1, P2RX2, P2RX3, P2RX4, P2RX5, P2RX6, P2RX7, PKD2, PKD2L1, PKD2L2, SCN10A, SCN11A, SCN1A, SCN2A, SCN3A, SCN4A, SCN5A, SCN8A, SCN9A, TPCN1, TPCN2, TRPA1, TRPC1, TRPC3, TRPC4, TRPC5, TRPC6, TRPC7, TRPM1, TRPM2, TRPM3, TRPM4, TRPM5, TRPM6, TRPM7, TRPM8, TRPV1, TRPV2, TRPV3, TRPV4, TRPV5, TRPV6, ZACN |
| Brain ion channel genes | CACNA1A, CACNA1B, CACNA1E, CACNA1G, CACNA1I, CHRNA2, CHRNA4, CHRNA6, CHRNA9, CHRNB2, CHRNB3, GABRA1, GABRA2, GABRA3, GABRA4, GABRA5, GABRA6, GABRB1, GABRB2, GABRB3, GABRD, GABRG1, GABRG2, GABRG3, GLRA1, GLRA2, GLRA3, GLRA4, GLRB, GRIA1, GRIA2, GRIA3, GRIA4, GRID1, GRID2, GRIK1, GRIK2, GRIK3, GRIK4, GRIK5, GRIN1, GRIN2A, GRIN2B, GRIN2C, GRIN2D, GRIN3A, HCN1, HCN2, HCN3, HCN4, HTR3B, KCNA1, KCNA2, KCNA4, KCNA6, KCNB1, KCNB2, KCNC1, KCNC2, KCNC3, KCND1, KCND2, KCND3, KCNF1, KCNG2, KCNG3, KCNH1, KCNH3, KCNH4, KCNH5, KCNH7, KCNH8, KCNJ2, KCNJ3, KCNJ4, KCNJ6, KCNJ9, KCNJ10, KCNJ12, KCNJ13, KCNK1, KCNK4, KCNK9, KCNK10, KCNK12, KCNK13, KCNK18, KCNN2, KCNN3, KCNQ2, KCNQ3, KCNQ5, KCNS1, KCNS2, KCNT1, KCNV1, P2RX7, PKD2L1, SCN1A, SCN2A, SCN3A, SCN8A, TRPC3, TRPC5, TRPC7, TRPM3 |
| Excitatory receptor genes | CHRNA1, CHRNA10, CHRNA2, CHRNA3, CHRNA4, CHRNA5, CHRNA6, CHRNA7, CHRNA9, CHRNB1, CHRNB2, CHRNB3, CHRNB4, CHRND, CHRNE, CHRNG, GRIA1, GRIA2, GRIA3, GRIA4, GRID1, GRID2, GRIK1, GRIK2, GRIK3, GRIK4, GRIK5, GRIN1, GRIN2A, GRIN2B, GRIN2C, GRIN2D, GRIN3A, GRIN3B. |
| NMDAR genes | ACTN2, ACTN4, ALDOC, ARF5, ATP1A1, ATP1A3, ATP1B1, ATP2B4, BAIAP2, CAMK2A, CAMK2B, CAPZA2, CAPZB, CDH2, CIT, CRMP1, CTNNB1, CYFIP1, DLG1, DLG2, DLG3, DLG4, DLGAP1, DLGAP2, DNM1, DPYSL2, FLNA, GAP43, GAPDH, GLUD1, GLUL, GNAS, GNB1, GNB2, GNB4, GRIN1, GSN, HOMER1, HSPA4, INA, IQSEC1, IQSEC2, MAP2, MAP2K1, MAPK3, MBP, MYO5A, NEFM, NSF, PKM, PLCB1, PLP1, PPP2R1A, PPP3CA, PRKAR2B, PRKCG, PTK2B, PTPN11, RAB3A, RAB6A, RAC1, SHANK1, SLC25A2, SLC25A3, SLC25A4, SNAP25, SPTBN1, SRC, STX1A, STXBP1, SYNGAP1, SYT1, TJP1, TUBA1B, YWHAE, YWHAG, YWHAH, YWHAZ. |
| PSD-95 genes | ABLIM1, ABR, ACAT1, ACO2, ACOT7, ADAM22, ALDOC, ANKS1A, ANKS1B, ARC, ARF3, ARPC4, ATP1B1, ATP5A1, ATP5B, ATP5C1, ATP5O, ATP6V0D1, BAIAP2, BTBD11, CACNG2, CAMK2A, CAMK2B, CAPZA2, CFL1, CLTC, CNP, CPNE4, CPNE7, CPNE8, DLG1, DLG2, DLG3, DLG4, DLGAP1, DLGAP2, DLGAP3, DLGAP4, DSTN, FAM81A, FRMPD3, FSCN1, GAPDH, GDA, GLUL, GNAO1, GPR123, GPX4, GRIA1, GRIA2, GRIA3, GRIA4, GRIK2, GRIK5, GRIN1, GRIN2A, GRIN2B, GRIN2D, IQSEC1, IQSEC2, KALRN, KCNA1, KCNA2, KCNA3, KCNA4, KCNAB1, KCNAB2, KCNJ10, KCNJ4, LGI1, MAPK1, MSRB2, NRXN1, NSF, PARK7, PCBP1, PDHA1, PDHB, PGAM5, PGK1, PHB2, PLP1, PPAP2B, PPP3CA, PPP3CB, PRDX1, PRDX2, PRRT1, RAC1, RPS14, RPS3, SDHA, SEPT11, SEPT5, SFXN3, SLC1A2, SLC25A4, SLC25A5, SLC4A4, SLC9A3R1, SPTBN1, STX1B, STXBP1, SUCLA2, SYN-GAP1, SYT1, TUBA1A, TUBA1C, TUBB2B, TUBB6, UBA52, VAMP2, VDAC1, VDAC2, VMAC, YWHAE, PKM, BEGAIN. |

| | |
|---|---|
| GO-GABAergic-Glutamatergic genes | ABHD6, ACAN, ADRA1A, ATP2B1, ATP2B2, ATP2B3, BSN, C1QBP, CALB1, CDH10, CLSTN1, CLSTN3, CNR1, CSPG5, CTBP1, CTBP2, DAG1, DRD1, DRD2, DRD3, ERBB4, ERC2, FLOT1, GABRD, GABRR1, GUCY1A3, KCND2, LRFN4, LRFN5, LRRTM1, LRRTM2, NLGN3, NLGN4X, NLGN4Y, NPTN, NRG1, NRXN1, PCDH17, PCLO, PHB2, PLCB1, PTPRO, SEPT11, SLC6A17, SLITRK1, SLITRK2, SV2A, SYT7. |
| GO-Glutamatergic genes | ABHD17A, ABHD17B, ABHD17C, ABLIM3, ABR, ACTB, ACTBL2, ACTC1, ACTN1, ACTN2, ADAM10, ADAM22, ADAM23, ADCY1, ADCY8, ADORA2A, ADORA2B, ADRA2C, AKAP9, AP2B1, AP3D1, APBA1, APOE, ARC, ARF1, ARF4, ARF6, ARHGAP22, ARHGAP39, ARHGAP44, ARHGEF2, ARPC2, ARPC5L, ATAD1, ATP2B4, BAI1, BAIAP2, BCAN, BCR, BZRAP1, CACNG2, CACNG3, CACNG4, CACNG5, CACNG7, CADPS, CADPS2, CALY, CAMK1, CAMK4, CAMKV, CANX, CBLN1, CBLN2, CDH1, CDH11, CDH8, CDK5, CDKL5, CHMP2B, CHRM1, CHRM2, CHRM3, CLSTN2, CNIH2, CNKSR2, CORO1A, CPLX1, CPLX2, CPT1C, CRTC1, CTNND1, CTTNBP2, DBN1, DGKB, DGKE, DGKI, DGKQ, DGKZ, DLG1, DLG2, DLG3, DLG4, DLGAP1, DLGAP2, DLGAP3, DLGAP4, DNAJB1, DNM1, DNM2, DNM3, DRD4, DRP2, DTNBP1, DVL1, DYNLL2, EFNB2, EFNB3, EGLN1, EIF4A3, EIF4E, EIF4EBP1, ELAVL1, EPHA4, EPHA7, EPHB1, EPHB2, EPS15, EPS8, ERBB2IP, FBXL20, FBXO45, FLNA, FLOT2, FLRT3, FRMPD4, FXYD6, FYN, FZD4, FZD9, GHRL, GHSR, GIPC1, GPC4, GPC6, GPM6A, GPR123, GRASP, GRIA1, GRID1, GRID2, GRIK2, GRIK3, GRIK5, GRIN1, GRIN2A, GRIN2C, GRIN3A, GRIP1, GRIPAP1, GRM1, GRM3, GSG1L, GSK3B, HIP1, HOMER1, HOMER2, HPCA, HRAS, HSPA8, HTR2A, HTR3A, IL1RAP, IL1RAPL1, IL1RAPL2, ITGB1, ITGB3, ITSN1, JAK2, KCNA1, KCNA3, KPNA1, KPNA2, LGI1, LIN7C, LPHN3, LPPR4, LRFN3, LRRC4, LRRC4B, LRRC4C, LRRK2, LRRTM3, LYN, MAPK10, MAPK14, MTOR, MYH10, MYO5A, NAPA, NCS1, NDRG1, NEDD4, NETO1, NLGN1, NOS1AP, NPTX1, NPTX2, NPTXR, NRCAM, NRG3, NRGN, NRP1, NRP2, NSG1, NTNG1, NTNG2, NTRK3, NUMB, OLFM2, OPHN1, P2RX1, P2RX6, P2RY1, P2RY4, PACSIN2, PAK2, PAK3, PCDH8, PFN1, PFN2, PIN1, PLAT, PLCB4, PLCG1, PLEKHA5, PLG, PORCN, PPFIA2, PPFIA3, PPM1H, PPP1CA, PPP1CC, PPP1R1B, PPP1R9A, PPP2R1A, PPP2R2A, PPP3CA, PPP3CB, PPP3CC, PRKAR1A, PRKAR1B, PRKAR2B, PRKCI, PRKCZ, PRR7, PRRT1, PRRT2, PSD2, PTK2B, PTPRD, PTPRS, PURA, RAB11A, RAB8A, RAC1, RAP1A, RELA, RGS14, RGS7BP, RGS9, RHOA, RNF19A, RNF216, RTN4R, SCN10A, SCN2A, SCRIB, SEMA3F, SH3GL1, SH3GL2, SH3GL3, SHANK1, SHISA6, SHISA7, SHISA9, SLC1A2, SLC1A6, SLC30A3, SLC4A8, SLITRK4, SNAP25, SNAP47, SORCS3, SPARC, SPARCL1, SPTBN1, SPTBN2, SRC, SRGN, STAT3, STAU1, STX1A, STX3, STX4, STXBP1, SYN3, SYNGAP1, SYNPO, SYT1, SYT4, SYT6, TIAM1, TNIK, TNR, UNC13A, UNC13B, USP46, USP8, VAMP1, VCP, VPS18, VPS35, WASF3, WNT3A, WNT5A, WNT7A, YES1, YWHAE, YWHAZ. |
| GO-GABAergic genes | CBLN4, CDH13, CNTN5, EFNA5, GABBR1, GABRA1, GABRA3, GABRA4, GABRA5, GABRB1, GABRB2, GABRB3, GABRG2, GABRG3, GABRR2, GAP43, GLRB, IGSF9B, IQSEC3, KCND3, MDGA1, NLGN2, SLC32A1, SLC6A1, SLC6A11, SLITRK3. |
| KEGG-Cholinergic genes | ACHE, ADCY1, ADCY2, ADCY3, ADCY4, ADCY5, ADCY6, ADCY7, ADCY8, ADCY9, AKT1, AKT2, AKT3, ATF4, BCL2, CACNA1A, CACNA1B, CACNA1C, CACNA1D, CACNA1F, CACNA1S, CAMK2A, CAMK2B, CAMK2D, CAMK2G, CAMK4, CHAT, CHRM1, CHRM2, CHRM3, CHRM4, CHRM5, CHRNA3, CHRNA4, CHRNA6, CHRNA7, CHRNB2, CHRNB4, CREB1, CREB3, CREB3L1, CREB3L2, CREB3L3, CREB3L4, CREB5, FOS, FYN, GNA11, GNAI1, GNAI2, GNAI3, GNAO1, GNAQ, GNB1, GNB2, GNB3, GNB4, GNB5, GNG10, GNG11, GNG12, GNG13, GNG2, GNG3, GNG4, GNG5, GNG7, GNG8, GNGT1, GNGT2, HRAS, ITPR1, ITPR2, ITPR3, JAK2, KCNJ12, KCNJ14, KCNJ2, KCNJ3, KCNJ4, KCNJ6, KCNQ1, KCNQ2, KCNQ3, KCNQ4, KCNQ5, KRAS, MAP2K1, MAPK1, MAPK3, NRAS, PIK3CA, PIK3CB, PIK3CD, PIK3CG, PIK3R1, PIK3R2, PIK3R3, PIK3R5, PIK3R6, PLCB1, PLCB2, PLCB3, PLCB4, PRKACA, PRKACB, PRKACG, PRKCA, PRKCB, PRKCG, SLC18A3, SLC5A7. |

| | |
|---|---|
| KEGG-Dopaminergic genes | ADCY5, AKT1, AKT2, AKT3, ARNTL, ARRB1, ARRB2, ATF2, ATF4, ATF6B, CACNA1A, CACNA1B, CACNA1C, CACNA1D, CALM1, CALM2, CALM3, CALML3, CALML4, CALML5, CALML6, CALY, CAMK2A, CAMK2B, CAMK2D, CAMK2G, CLOCK, COMT, CREB1, CREB3, CREB3L1, CREB3L2, CREB3L3, CREB3L4, CREB5, DDC, DRD1, DRD2, DRD3, DRD4, DRD5, FOS, GNAI1, GNAI2, GNAI3, GNAL, GNAO1, GNAQ, GNAS, GNB1, GNB2, GNB3, GNB4, GNB5, GNG10, GNG11, GNG12, GNG13, GNG2, GNG3, GNG4, GNG5, GNG7, GNG8, GNGT1, GNGT2, GRIA1, GRIA2, GRIA3, GRIA4, GRIN2A, GRIN2B, GSK3A, GSK3B, ITPR1, ITPR2, ITPR3, KCNJ3, KCNJ5, KCNJ6, KCNJ9, KIF5A, KIF5B, KIF5C, LRTOMT, MAOA, MAOB, MAPK10, MAPK11, MAPK12, MAPK13, MAPK14, MAPK8, MAPK9, PLCB1, PLCB2, PLCB3, PLCB4, PPP1CA, PPP1CB, PPP1CC, PPP1R1B, PPP2CA, PPP2CB, PPP2R1A, PPP2R1B, PPP2R2A, PPP2R2B, PPP2R2C, PPP2R2D, PPP2R3A, PPP2R3B, PPP2R3C, PPP2R5A, PPP2R5B, PPP2R5C, PPP2R5D, PPP2R5E, PPP3CA, PPP3CB, PPP3CC, PRKACA, PRKACB, PRKACG, PRKCA, PRKCB, PRKCG, SCN1A, SLC18A1, SLC18A2, SLC6A3, TH. |
| KEGG-mTOR genes | AKT1, AKT1S1, AKT2, AKT3, ATP6V1A, ATP6V1B1, ATP6V1B2, ATP6V1C1, ATP6V1C2, ATP6V1D, ATP6V1E1, ATP6V1E2, ATP6V1F, ATP6V1G1, ATP6V1G2, ATP6V1G3, ATP6V1H, BRAF, CAB39, CAB39L, CHUK, CLIP1, DDIT4, DEPDC5, DEPTOR, DVL1, DVL2, DVL3, EIF4B, EIF4E, EIF4E1B, EIF4E2, EIF4EBP1, FLCN, FNIP1, FNIP2, FZD1, FZD10, FZD2, FZD3, FZD4, FZD5, FZD6, FZD7, FZD8, FZD9, GATSL1, GATSL3, GRB10, GRB2, GSK3B, HRAS, IGF1, IGF1R, IKBKB, INS, INSR, IRS1, KRAS, LAMTOR1, LAMTOR2, LAMTOR3, LAMTOR4, LAMTOR5, LPIN1, LPIN2, LPIN3, LRP5, LRP6, MAP2K1, MAP2K2, MAPK1, MAPK3, MAPKAP1, MIOS, MLST8, MTOR, NPRL2, NPRL3, NRAS, PDPK1, PIK3CA, PIK3CB, PIK3CD, PIK3R1, PIK3R2, PIK3R3, PRKAA1, PRKAA2, PRKCA, PRKCB, PRKCG, PRR5, PTEN, RAF1, RHEB, RHOA, RICTOR, RNF152, RPS6, RPS6KA1, RPS6KA2, RPS6KA3, RPS6KA6, RPS6KB1, RPS6KB2, RPTOR, RRAGA, RRAGB, RRAGC, RRAGD, SEC13, SEH1L, SESN2, SGK1, SKP2, SLC38A9, SLC3A2, SLC7A5, SOS1, SOS2, STK11, STRADA, STRADB, TBC1D7, TELO2, TNF, TNFRSF1A, TSC1, TSC2, TTI1, ULK1, ULK2, WDR24, WDR59, WNT1, WNT10A, WNT10B, WNT11, WNT16, WNT2, WNT2B, WNT3, WNT3A, WNT4, WNT5A, WNT5B, WNT6, WNT7A, WNT7B, WNT8A, WNT8B, WNT9A, WNT9B |
| KEGG-GABAergic genes | ABAT, ADCY1, ADCY2, ADCY3, ADCY4, ADCY5, ADCY6, ADCY7, ADCY8, ADCY9, CACNA1A, CACNA1B, CACNA1C, CACNA1D, CACNA1F, CACNA1S, GABARAP, GABARAPL1, GABARAPL2, GABBR1, GABBR2, GABRA1, GABRA2, GABRA3, GABRA4, GABRA5, GABRA6, GABRB1, GABRB2, GABRB3, GABRD, GABRE, GABRG1, GABRG2, GABRG3, GABRP, GABRQ, GABRR1, GABRR2, GAD1, GAD2, GLS, GLS2, GLUL, GNAI1, GNAI2, GNAI3, GNAO1, GNB1, GNB2, GNB3, GNB4, GNB5, GNG10, GNG11, GNG12, GNG13, GNG2, GNG3, GNG4, GNG5, GNG7, GNG8, GNGT1, GNGT2, GPHN, HAP1, KCNJ6, NSF, PLCL1, PRKACA, PRKACB, PRKACG, PRKCA, PRKCB, PRKCG, SLC12A5, SLC32A1, SLC38A1, SLC38A2, SLC38A5, SLC6A1, SLC6A11, SLC6A12, SLC6A13, SRC, TRAK2 |
| KEGG-GABAergic-Glutamatergic genes | ADCY1, ADCY2, ADCY3, ADCY4, ADCY5, ADCY6, ADCY7, ADCY8, ADCY9, CACNA1A, CACNA1C, CACNA1D, GLS, GLS2, GLUL, GNAI1, GNAI2, GNAI3, GNAO1, GNB1, GNB2, GNB3, GNB4, GNB5, GNG10, GNG11, GNG12, GNG13, GNG2, GNG3, GNG4, GNG5, GNG7, GNG8, GNGT1, GNGT2, PRKACA, PRKACB, PRKACG, PRKCA, PRKCB, PRKCG, SLC38A1, SLC38A2. |
| KEGG-Glutamatergic genes | ADCY1, ADCY2, ADCY3, ADCY4, ADCY5, ADCY6, ADCY7, ADCY8, ADCY9, ADRBK1, ADRBK2, CACNA1A, CACNA1C, CACNA1D, DLG4, DLGAP1, GLS, GLS2, GLUL, GNAI1, GNAI2, GNAI3, GNAO1, GNAQ, GNAS, GNB1, GNB2, GNB3, GNB4, GNB5, GNG10, GNG11, GNG12, GNG13, GNG2, GNG3, GNG4, GNG5, GNG7, GNG8, GNGT1, GNGT2, GRIA1, GRIA2, GRIA3, GRIA4, GRIK1, GRIK2, GRIK3, GRIK4, GRIK5, GRIN1, GRIN2A, GRIN2B, GRIN2C, GRIN2D, GRIN3A, GRIN3B, GRM1, GRM2, GRM3, GRM4, GRM5, GRM6, GRM7, GRM8, HOMER1, HOMER2, HOMER3, ITPR1, ITPR2, ITPR3, JMJD7-PLA2G4B, KCNJ3, MAPK1, MAPK3, PLA2G4A, PLA2G4B, PLA2G4C, PLA2G4D, PLA2G4E, PLA2G4F, PLCB1, PLCB2, PLCB3, PLCB4, PLD1, PLD2, PPP3CA, PPP3CB, PPP3CC, PPP3R1, PPP3R2, PRKACA, PRKACB, PRKACG, PRKCA, PRKCB, PRKCG, SHANK1, SHANK2, SHANK3, SLC17A6, SLC17A7, SLC17A8, SLC1A1, SLC1A2, SLC1A3, SLC1A6, SLC1A7, SLC38A1, SLC38A2, TRPC1. |

| | |
|---|---|
| KEGG-Synaptic vesicle genes | AP2A1, AP2A2, AP2B1, AP2M1, AP2S1, ATP6V0A1, ATP6V0A2, ATP6V0A4, ATP6V0B, ATP6V0C, ATP6V0D1, ATP6V0D2, ATP6V0E1, ATP6V0E2, ATP6V1A, ATP6V1B1, ATP6V1B2, ATP6V1C1, ATP6V1C2, ATP6V1D, ATP6V1E1, ATP6V1E2, ATP6V1F, ATP6V1G1, ATP6V1G2, ATP6V1G3, ATP6V1H, CACNA1A, CACNA1B, CLTA, CLTB, CLTC, CLTCL1, CPLX1, CPLX2, CPLX3, CPLX4, DNM1, DNM2, DNM3, NAPA, NSF, RAB3A, RIMS1, SLC17A6, SLC17A7, SLC17A8, SLC18A1, SLC18A2, SLC18A3, SLC1A1, SLC1A2, SLC1A3, SLC1A6, SLC1A7, SLC32A1, SLC6A1, SLC6A11, SLC6A12, SLC6A13, SLC6A2, SLC6A3, SLC6A4, SLC6A5, SLC6A7, SLC6A9, SNAP25, STX1A, STX1B, STX2, STX3, STXBP1, SYT1, TCIRG1, UNC13A, UNC13B, UNC13C, VAMP2. |
| Reactome-GABAergic genes | ABAT, ADCY1, ADCY2, ADCY3, ADCY4, ADCY5, ADCY6, ADCY7, ADCY8, ADCY9, ALDH5A1, ARHGEF9, CPLX1, DNAJC5, GABBR1, GABBR2, GABRA1, GABRA2, GABRA3, GABRA4, GABRA5, GABRA6, GABRB1, GABRB2, GABRB3, GABRG2, GABRG3, GABRQ, GABRR1, GABRR2, GAD1, GAD2, GNAI1, GNAI2, GNAI3, GNAL, GNAT3, GNB1, GNB2, GNB3, GNB4, GNB5, GNG10, GNG11, GNG12, GNG13, GNG2, GNG3, GNG4, GNG5, GNG7, GNG8, GNGT1, GNGT2, HSPA8, KCNJ10, KCNJ12, KCNJ15, KCNJ16, KCNJ2, KCNJ3, KCNJ4, KCNJ5, KCNJ6, KCNJ9, NPTN, RAB3A, RIMS1, SLC32A1, SLC6A1, SLC6A11, SLC6A12, SLC6A13, SNAP25, STX1A, STXBP1, SYT1, VAMP2. |
| Reactome-Glutamatergic genes | ACTN2, ADCY1, ADCY8, AKAP5, AP2A1, AP2A2, AP2B1, AP2M1, AP2S1, APBA1, ARHGEF7, ARL6IP5, BZRAP1, CACNG2, CACNG3, CACNG4, CACNG8, CALM1, CAMK1, CAMK2A, CAMK2B, CAMK2D, CAMK2G, CAMK4, CAMKK1, CAMKK2, CASK, CPLX1, CREB1, DLG1, DLG2, DLG3, DLG4, EPB41L1, ERBB4, GIT1, GLS, GLS2, GLUL, GNB1, GNB2, GNB3, GNB4, GNB5, GNG10, GNG11, GNG12, GNG13, GNG2, GNG3, GNG4, GNG5, GNG7, GNG8, GNGT1, GNGT2, GRIA1, GRIA2, GRIA3, GRIA4, GRIK1, GRIK2, GRIK3, GRIK4, GRIK5, GRIN1, GRIN2A, GRIN2B, GRIN2C, GRIN2D, GRIN3A, GRIN3B, GRIP1, HRAS, KIF17, KPNA2, KRAS, LIN7A, LIN7B, LIN7C, LRRC7, MAPK1, MAPK3, MAPT, MDM2, MYO6, NBEA, NCALD, NRAS, NRG1, NRGN, NSF, PDPK1, PICK1, PLCB1, PLCB2, PLCB3, PPFIA1, PPFIA2, PPFIA3, PPFIA4, PPM1E, PPM1F, PRKAA1, PRKAA2, PRKAB1, PRKAB2, PRKACA, PRKACB, PRKACG, PRKAG1, PRKAG2, PRKAG3, PRKAR1A, PRKAR1B, PRKAR2A, PRKAR2B, PRKCA, PRKCB, PRKCG, PRKX, RAB3A, RAC1, RASGRF1, RASGRF2, RIMS1, RP11-683L23.1, RPS6KA1, RPS6KA2, RPS6KA3, RPS6KA6, SLC17A7, SLC1A1, SLC1A2, SLC1A3, SLC1A6, SLC1A7, SLC38A1, SLC38A2, SNAP25, SRC, STX1A, STXBP1, SYT1, TSPAN7, TUBA1A, TUBA1B, TUBA1C, TUBA3C, TUBA3D, TUBA3E, TUBA4A, TUBA8, TUBAL3, TUBB1, TUBB2A, TUBB2B, TUBB3, TUBB4A, TUBB4B, TUBB6, TUBB8, UNC13B, VAMP2. |
| Reactome-NeurXG genes | APBA1, APBA2, APBA3, BEGAIN, CASK, DBNL, DLG2, DLG3, DLG4, DLGAP1, DLGAP2, DLGAP3, DLGAP4, EPB41, EPB41L1, EPB41L2, EPB41L3, EPB41L5, GRIN1, GRIN2A, GRIN2B, GRIN2C, GRIN2D, GRM1, GRM5, HOMER1, HOMER2, HOMER3, LIN7A, LIN7B, LIN7C, LRRTM1, LRRTM2, LRRTM3, LRRTM4, NLGN1, NLGN2, NLGN3, NLGN4X, NLGN4Y, NRXN1, NRXN2, NRXN3, PDLIM5, SHANK1, SHANK2, SHANK3, SHARPIN, SIPA1L1, STX1A, STXBP1, SYT1, SYT10, SYT12, SYT2, SYT7, SYT9. |
| Reactome-PresynDep genes | CACNA1A, CACNA1B, CACNA1E, CACNA2D1, CACNA2D2, CACNA2D3, CACNB1, CACNB2, CACNB3, CACNB4, CACNG2, CACNG4. |
| Reactome-RTP genes | IL1RAP, IL1RAPL1, IL1RAPL2, LRRC4B, NTRK3, PPFIA1, PPFIA2, PPFIA3, PPFIA4, PPFIBP1, PPFIBP2, PTPRD, PTPRF, PTPRS, SLITRK1, SLITRK2, SLITRK3, SLITRK4, SLITRK5, SLITRK6. |
| Reactome-SynAdhesion genes | DLG1, DLG3, DLG4, FLOT1, FLOT2, GRIA1, GRIA3, GRIA4, GRIN1, GRIN2A, GRIN2B, GRIN2C, GRIN2D, LRFN1, LRFN2, LRFN3, LRFN4, PTPRD, PTPRF, PTPRS, RTN3. |
| Dominant epilepsy genes | ALG13, CDKL5, CHD2, CHRNA2, CHRNA4, CHRNB2, DEPDC5, DNM1, EEF1A2, GABRA1, GABRB3, GABRG2, GNAO1, GRIN1, GRIN2A, GRIN2B, HCN1, HNRNPU, KCNA2, KCNB1, KCNC1, KCNMA1, KCNQ2, KCNQ3, KCNT1, LGI1, MEF2C, PCDH19, PRICKLE2, PRRT2, SCN1A, SCN1B, SCN2A, SCN8A, SCN9A, SIK1, SLC2A1, SLC35A2, SLC6A1, SPTAN1, STX1B, STXBP1, SYNGAP1 |

| | |
|---|---|
| GWAS genes | SCN1A, TTC21B, TNKS, PABPC4, KIAA0408, IL6, CAMTA1, IQSEC1, VRK2, USP45, BMP8A, MACF1, FLJ00104, POU6F2, AC010536.1, PNISR, PXN, RPLP0, FAM69A, ZKSCAN3, KLHDC4, ZSCAN31, FAXC, IPO7, CNEP1R1, PRPSAP2, ZNF512B, SLCO3A1, CDPF1, PPARA, WIPF1, SOGA3, FBXW10, UCKL1, GCN1L1, APOH, TTC38, BPTF, CADPS2, SCN9A, ITPKB, PRPF6, TVP23B, SBDS, HEATR3, CKAP4, RP11-1167A19.2, SAMD10, MRS2, PKDREJ, JAM2, CDC42EP4, PRDM13, C16orf96, KATNAL1, HEYL, ZSCAN12, NOB1, SERTAD4, KIAA0754, THSD7B, NT5DC4, CD46, COQ3, TCP11L2, ZNF143, CTSZ, FAM8A1, ILF3, CPLX2, DGKH, SLC44A2, MAGI2, TCEA2, FANCL, ANKRD27, PAX9, AC007421.1, DIAPH2, TUBB1, SOX15, ANO8, AC011475.1, GTSE1, RPL5, KLF1, CRTAC1, TMEM182, ZMIZ1, ASPM, ZSCAN23, FEZ1, TUBB2A, GCDH, C9orf64, ADORA2B, FAM122A, FLRT2, DNASE2, SH2B1 |
| GWAS-focal genes | SCN1A, IL6, TNKS, KIAA0408, ANKRD27, AC010536.1, TTC21B, FLJ00104, RGS9BP, TFPI, SLCO3A1, CKAP4, FUNDC1, KCNMA1, USP45, PABPC4, LOXHD1, FAXC, KLHDC4, TCP11L2, LHX5, CUL4A, UPB1, SERPINE2, NUDT19, ASPM, CILP, GOT1, CYSLTR2, SOGA3, PNISR, SKOR2, GGT1, PDE8B, PXN, PCID2, SNRPD3, PAPD5, TNS3, PPARA, GUCD1, POM121L2, WIPF1, OR4F6, TSPAN31, RPL9, B4GALNT1, AGAP2, OR4F15, CDPF1, TTC38, PARP16, NT5DC4, CABLES2, AGAP2-AS1, OS9, HNRNPH3, ERVV-1, FAM151B, AVIL, TUBB2A, SCN9A, BCAR3, NCAN, PBLD, PRPF38B, MARCH9, FGF7, VEGFA, IQSEC1, PKDREJ, NDUFA10, SLC13A4, LIAS, KLF1, PRLR, VWA2, CCL23, FAM227B, ZBTB41, BPHL, LIPF, MNT, TSPAN8, TXN2, BPTF, COQ3, A2ML1, HLF, ZFYVE16, TDRD1, TSFM, CTDSP2, CYTH4, POTEE, BMP8A, METTL21B, CDK4, SDS, CCL16. |
| GWAS-GGE genes | VRK2, SCN1A, AP3D1, DNAH14, SPOCK1, FANCL, KRTAP8-1, PRR15L, FAM49B, CDK5RAP3, CAMTA1, C1orf134, RSG1, PPP2R2B, C9orf64, SKAP1, SETD1A, AC135048.1, TMEM115, FBXL19, MOB3A, STX1B, FADS2, ORAI3, LSM12, HSD3B7, ATXN7L3, HDAC5, TMEM182, ZNF646, ZNF668, NOL4, FBXO42, SH3BP1, RMI1, ZNF143, CD47, LGALS1, NFE2L1, STX4, ZNF668, PCDH7, MYCBPAP, NOL12, COPZ2, YIF1A, LDLRAD3, IPO7, ZNF226, TVP23B, NPRL2, HNRNPK, ZNF512B, PDXP, FADS3, IZUMO4, BCKDK, FADS1, INHBA, G6PC3, SP6, MFSD9, RAPGEF2, HHLA2, ZMYND10, PRSS53, UCKL1, MFAP3, XXcos-LUCA11.5, PRPF6, RASSF1, TUSC2, KIF27, ACVRL1, HYAL2, IL20RB, CYB561D2, RP11-196G11.1, TOMM40, UBTF, TM6SF1, BCL7C, STMND1, DOT1L, RIMS1, SCRN2, JMY, HYAL1, ACVR1B, SAMD10, C17orf53, TMEM107, PER1, ZNF91, CTPS1, TMEM41B, OXCT1, C17orf59, MRPL33, PRPSAP2. |
| Genetic epilepsy genes | ALDH7A1, CACNA1A, CACNA1H, CACNB4, CASR, CNTN2, EFHC1, EPM2A, GABRA1, GABRB3, GABRD, GABRG2, GPHN, KCNA2, KCNC1, KCNMA1, NIPA2, NRXN1, PCDH7, PLCB1, RBFOX1, RORB, SCN1A, SCN1B, SCN9A, SLC2A1, STX1B, TBC1D24. |
| Focal epilepsy genes | CHRNA2, CHRNA4, CHRNB2, CPA6, DEPDC5, GRIN2A, GRIN2B, KCNA1, KCNQ2, KCNQ3, KCNT1, LGI1, NPRL2, NPRL3, PRRT2, RBFOX1, RBFOX3, SCN2A, SCN8A, TBC1D24. |
| Developmental and Epileptic Encephalopathies genes | ALG13, ARHGEF9, ARX, CACNA1A, CASK, CDKL5, CHD2, DNM1, EEF1A2, FOXG1, GABRA1, GABRB3, GNAO1, GNB1, GPHN, GRIN1, GRIN2A, GRIN2B, GRIN2D, HCN1, IQSEC2, KCNA2, KCNB1, KCNQ2, KCNT1, MBD5, MECP2, MEF2C, PCDH19, PIGA, PURA, SCN1A, SCN2A, SCN8A, SIK1, SLC1A2, SLC2A1, SLC35A2, SLC6A1, SLC6A8, SLC9A6, SPTAN1, STXBP1, SYN1, SYNGAP1, TSC1, TSC2, UBE3A, WDR45, ZEB2. |

| | |
|---|---|
| FMPR genes | AAK1, AATK, ABCA2, ABCA3, ABCG1, ABR, ACLY, ACO2, ACTB, ADAP1, ADARB1, ADCY1, ADCY5, ADD1, ADNP, ADRBK1, AFF3, AFF4, AGAP1, AGAP2, AGAP2-AS1, AGAP3, AGO1, AGO2, AGPAT3, AGRN, AGTPBP1, AHDC1, AKAP6, AKAP9, AKT3, ALDOA, ALDOC, ALS2, AMPH, ANAPC1, ANK1, ANK2, ANK3, ANKRD11, ANKRD17, ANKRD52, AP1B1, AP2A1, AP2A2, AP2B1, AP3D1, APBA1, APBB1, APC, APC2, APLP1, APOE, APP, ARAP2, AREL1, ARF3, ARFGEF1, ARHGAP20, ARHGAP21, ARHGAP23, ARHGAP32, ARHGAP33, ARHGAP35, ARHGAP44, ARHGEF11, ARHGEF12, ARHGEF17, ARHGEF2, ARHGEF4, ARHGEF7, ARID1A, ARID1B, ARID2, ARNT2, ARPP21, ARRB1, ARVCF, ASH1L, ATF7IP, ATG2A, ATG2B, ATG9A, ATMIN, ATN1, ATP13A2, ATP1A1, ATP1A2, ATP1A3, ATP1B1, ATP2A2, ATP2B2, ATP2B4, ATP5A1, ATP5B, ATP6V0A1, ATP6V0D1, ATP6V1B2, ATP9A, ATXN1, AUTS2, B3GAT1, BAG6, BAI1, BAI2, BAP1, BAZ2A, BCAN, BCL9L, BCR, BIRC6, BMPR2, BPTF, BRD4, BRINP1, BRINP2, BRSK1, BRSK2, BSN, BZRAP1, C19orf26, C2CD2L, CABIN1, CACNA1A, CACNA1B, CACNA1E, CACNA1G, CACNA1I, CACNB1, CACNB3, CADPS, CALM1, CALM3, CAMK2A, CAMK2B, CAMK2N1, CAMKK2, CAMSAP1, CAMSAP2, CAMTA1, CAMTA2, CAND1, CASKIN1, CBX6, CCSER2, CDC42BPA, CDC42BPB, CDK16, CDK17, CDK5R1, CDK5R2, CDKL5, CELF5, CELSR2, CELSR3, CEP170B, CHD3, CHD4, CHD5, CHD6, CHD8, CHN1, CHN2, CHST2, CIC, CIT, CKAP5, CKB, CLCN3, CLEC16A, CLIP3, CLSTN1, CLTC, CLUH, CNP, COBL, COPG1, CPE, CPLX1, CPLX2, CPT1C, CREBBP, CRMP1, CRTC1, CTBP1, CTNNB1, CTNND2, CUL9, CUX1, CUX2, DAB2IP, DAGLA, DAPK1, DCAF6, DCLK1, DCTN1, DDN, DDX24, DENND5A, DGCR2, DGKZ, DHX30, DICER1, DIDO1, DIP2A, DIP2B, DIP2C, DIRAS2, DISP2, DLC1, DLG2, DLG4, DLG5, DLGAP1, DLGAP2, DLGAP3, DLGAP4, DMTN, DMWD, DMXL2, DNAJC6, DNM1, DOCK3, DOCK4, DOCK9, DOPEY1, DOPEY2, DOT1L, DPP8, DPYSL2, DSCAM, DSCAML1, DST, DTNA, DTX1, DUSP8, DYNC1H1, DZANK1, EEF1A2, EEF2, EGR1, EHMT1, EHMT2, EIF4G1, EIF4G2, EIF4G3, ELFN2, ELMO2, EMC1, EML2, ENC1, EP300, EP400, EPB41L1, EPHA4, EPN1, EXTL3, FAM115A, FAM120A, FAM160A2, FAM171B, FAM179B, FAM21A, FAM65A, FAM91A1, FASN, FAT1, FAT2, FAT4, FBXL16, FBXL19, FBXO41, FCHO1, FKBP8, FOXK2, FOXO3, FRMPD4, FRY, FSCN1, FYN, GABBR1, GABBR2, GARNL3, GAS7, GBF1, GCN1L1, GIT1, GLUL, GNAL, GNAO1, GNAS, GNAZ, GNB1, GPAM, GPM6A, GPR158, GPR162, GPRIN1, GRAMD1B, GRIK3, GRIK5, GRIN1, GRIN2A, GRIN2B, GRM4, GRM5, GSK3B, GTF3C1, GTF3C2, HCFC1, HCN2, HDAC4, HDAC5, HDLBP, HEATR5B, HERC1, HERC2, HIPK1, HIPK3, HIVEP1, HIVEP2, HIVEP3, HK1, HMGXB3, HNRNPUL1, HSP90AB1, HTT, HUWE1, IDS, IGSF9B, INPP4A, INTS1, IPO13, IPO4, IPO5, IQSEC2, IQSEC3, IRF2BPL, IRS2, ITPR1, ITSN1, JAK1, JPH3, JPH4, KALRN, KAT6A, KCNA2, KCNB1, KCNC3, KCND2, KCNH1, KCNH3, KCNH7, KCNMA1, KCNQ2, KCNQ3, KCNT1, KDM4B, KDM5C, KDM6B, KIAA0100, KIAA0226, KIAA0430, KIAA0947, KIAA1045, KIAA1109, KIAA1244, KIAA1549L, KIAA2018, KIF1A, KIF1B, KIF21A, KIF21B, KIF3C, KIF5A, KIFC2, KLC1, KLHL22, KMT2A, KMT2C, KMT2D, KMT2E, KNDC1, LARGE, LARS2, LHFPL4, LINGO1, LLGL1, LMTK2, LMTK3, LPHN1, LPHN3, LPIN2, LPPR4, LRP1, LRP3, LRP8, LRRC41, LRRC4B, LRRC7, LRRC8B, LRRN2, LYNX1, LZTS3, MACF1, MADD, MAGED1, MAGI2, MAN2A2, MAP1A, MAP1B, MAP2, MAP3K12, MAP4, MAP4K4, MAP7D1, MAPK1, MAPK4, MAPK8IP1, MAPK8IP3, MAPKBP1, MAST1, MAST2, MAST4, MAZ, MBD5, MBP, MED13, MED13L, MED14, MED16, MEF2D, MFHAS1, MGAT5B, MIB1, MICAL2, MINK1, MKL2, MMP24, MON2, MPRIP, MTMR4, MTOR, MTSS1L, MYH10, MYO10, MYO16, MYO18A, MYO5A, MYT1L, NACAD, NAT8L, NAV1, NAV2, NAV3, NBEA, NCAN, NCDN, NCKAP1, NCOA1, NCOA2, NCOA6, NCOR1, NCOR2, NCS1, NDRG2, NDRG4, NDST1, NEDD4, NEURL4, NF1, NFIC, NFIX, NGEF, NHSL1, NISCH, NLGN2, NLGN3, NOMO1, NPAS2, NPTXR, NR2F1, NRGN, NRIP1, NRXN1, NRXN2, NRXN3, NSD1, NSF, NSMF, NTRK2, NTRK3, NUP98, NWD1, OGDH, OLFM1, OXR1, PACS1, PACS2, PAK6, PCDH1, PCDH10, PCDH7, PCDH9, PCDHA4, PCDHAC2, PCDHGA12, PCDHGC3, PCLO, PCNX, PCNXL2, PCNXL3, PDE2A, PDE4B, PDE4DIP, PDE8B, PDS5B, PDZD2, PDZD8, PEG3, PER1, PFKM, PGM2L1, PHF12, PHF20, PHLDB1, PHYHIP, PI4KA, PIGQ, PIKFYVE, PINK1, PIP5K1C, PITPNM1, PITPNM2, PJA2, PLAA, PLEKHM2, PLK3, PLPPR5, PLEC, PLXNA2, POMC, POPDC2, POT1, PPARGC1A, PRC1, PRDM9, PRKAR1B, PRKCE, PRKCI, PRKCZ, PRMT2, PRMT3, PRMT5, PSEN1, PSEN2, PTPN6, PTPN11, PTPN12, PTPN14, PTPN18, PTPN21, PTCH1, PTCHD1, PTGDR2, PTPRT, PUM1, PXDNL, QKI |

| | |
|---|---|
| FMPR genes | RAB7A, RAB9B, RAB11FIP4, RAB31, RAB40A, RAB3B, RAB5A, RAB5B, RAB9A, RAP1GAP, RAPGEF1, RAPGEF2, RASSF5, RBM3, RBM4, RBM5, RBM8A, RCBTB2, RCN1, REXO1, RHOG, RHOT1, RGS6, RGS7, RGS9, RHPN2, RIMKLB, RIT1, RLIM, RNF10, RNF11, RNF123, RNF43, ROBO1, ROBO2, RPA1, RPL7, RPL9, RPL10, RPL11, RPS11, RPS12, RPS3, RPS7, RPS8, RPS9, RPS10, RPS13, RPS14, RPS15, RPS16, RPS19, RPS21, RPS24, RPS26, RPS27, RPS27A, RPS28, RPS29, RPS30, RPS33, RPS34, RPSA, RTN1, RUFY1, RUFY3, S100A10, S100A11, S100A16, S100A1, S100B, S100G, S100P, SACS, SART1, SCAMP5, SCGB3A2, SCN1A, SCN2A, SCN3A, SCN4A, SCN8A, SDC4, SEMA4A, SEMA6A, SEMA6D, SERTAD1, SERPINB10, SETD5, SETX, SF3B1, SF3B3, SFMBT1, SHANK2, SHANK3, SHC1, SHOC2, SHPRH, SLC12A6, SLC16A1, SLC17A4, SLC22A1, SLC25A13, SLC26A2, SLC27A1, SLC29A4, SLC30A1, SLC33A1, SLC35A2, SLC36A4, SLC39A12, SLC41A2, SLC4A2, SLC5A2, SLC6A1, SLC6A2, SLC7A5, SLC9A1, SLC9A3, SLC9A4, SLC9A5, SMARCA4, SMARCB1, SMARCE1, SMC1A, SMC2, SMO, SND1, SNX2, SNX27, SNX30, SNX5, SNX6, SP3, SPATA2, SPECC1, SPG7, SPG21, SPRY2, SRGAP3, SRM, SRSF11, SRSF2, SSB, STAM2, STARD10, STK17A, STK11, STK39, STOML3, STRA6, STRN, SUGP2, SULT1A1, SULT1A2, SULT1A3, SULT1B1, SULT2A1, SULT2B1, SUMO1, SUMO2, SUMO3, SVIL, SYBU, SYNE2, SYNE3, SYNCRIP, SYT1, SYT2, SYT4, SYT7, TAF1, TAF5, TAF6, TAF7, TAF9, TANGO1, TAPT1, TAPT2, TBC1D20, TBC1D25, TBC1D4, TBX1, TBX3, TBX5, TCF3, TCF7, TCF7L1, TCF7L2, TCHH, TDP1, TEAD2, TET1, TET2, TFAP2A, TFAP2B, TFAP2C, TGIF1, TGIF2, THAP1, THAP2, TIA1, TLR4, TLE1, TLE4, TLK1, TMEM106B, TMEM14A, TMEM16A, TMEM16B, TMEM16C, TMEM16D, TMEM16E, TMEM16F, TMEM16G, TMEM16H, TMEM16J, TMEM16K, TMEM16L, TMEM16P, TMEM192, TMEM30A, TMEM38A, TMEM48, TMEM59L, TMEM9B, TMEM9C, TNNI2, TNNI3, TNS1, TOP2A, TOP2B, TP63, TRAPPC9, TRIM2, TRIM3, TRIM5, TRIM9, TRPC4, TRPC6, TRPS1, TSC1, TSC2, TSGA10, TSPEAR, TSPAN12, TSPAN8, TTF1, TTC21A, TUBB, TUBGCP2, TUBGCP5, TUBT, TYRO3, UBE2B, UBE2D1, UBE2D2, UBE2D3, UBE2E1, UBE2E2, UBE2G2, UBE2H, UBE2I, UBE2J2, UBE2L3, UBE2L6, UBE2N, UBE2V1, UBE2V2, UBQLN1, UBQLN2, UBQLN4, UCN, UGDH, UHRF1, UHRF2, UNC13A, UNC13B, UNC5C, UNC80, UPF3B, UPK3A, USP1, USP2, USP3, USP4, USP5, USP7, USP9X, VAMP1, VAMP2, VAX1, VCP, VIM, VIPR1, VIPR2, VPS13C, VPS13D, VPS13A, VPS35, VPS26A, VPS26B, WDR4, WDR45, WDR45B, WDR5, WDR62, WDR66, WDR68, WDR76, WDR78, WDR80, WDR81, WDR82, WDR9, XBP1, XCL1, XPO1, XPR1, YARS, YWHAE, YWHAZ |
| NDD-E genes | ALG13, ARHGEF9, ARID1B, ASXL3, CDKL5, CHD2, COL4A3BP, DNM1, DYRK1A, EEF1A2, FOXG1, GABRB2, GABRB3, GNAO1, GRIN2A, GRIN2B, HNRNPU, KCNH1, KCNQ2, KIAA2022, MECP2, MEF2C, PURA, SCN1A, SCN2A, SCN8A, SLC35A2, SLC6A1, SMC1A, SNAP25, STXBP1, SYNGAP1, WDR45 |
| MGI genes | ACHE, ACP2, ADAM22, ADAMTS4, ADARB1, AIFM1, AKT3, ALDH5A1, ALPL, AMPH, ANK3, AP3B2, AP3M2, APC2, ARX, ASPA, ATAD1, ATCAY, ATOX1, ATP1A3, ATP7A, ATRN, AVPR1B, BCKDK, BHLHE40, BLOC1S6, BMI1, BRAF, BSN, C1QA, CA7, CACNA1A, CACNA2D2, CACNB4, CACNG2, CAMK2A, CD3E, CDK5R1, CDKN2A, CELF4, CERS1, CHN1, CHRNA4, CHRNA7, CIT, CLCN3, CLCN7, CLN3, CLN6, CLN8, CNP, CNR1, CNTN2, CNTN5, CNTNAP2, COL2A1, COMMD3-BMI1, CPLX1, CREBBP, CSTB, CTSD, DBNL, DGKD, DGKE, DMTF1, DNM1, DRD2, DSCAM, DST, EFHC1, ELAVL3, FAIM2, FCGR2B, FGF14, FGF7, FMR1, FOSB, FYN, FZD9, G6PC, GABBR1, GABBR2, GABRA1, GABRB3, GABRD, GABRG2, GAD2, GAL, GAN, GDI1, GFAP, GLRA1, GLRA3, GNAO1, GNG3, GPR98, GRIA2, GRIA4, GRIK2, GRIK5, GRIN2A, GRM5, GRM7, HCN2, HCRTR1, HDAC4, HELT, HEXA, HEXB, HPRT1, HRH2, HTR1A, HTR2C, HTR4, HTR7, HTRA2, HTT, IL6, IMPA1, ITPR1, KCNA1, KCNA2, KCNA4, KCNAB2, KCNC2, KCNH3, KCNJ11, KCNJ6, KCNK2, KCNMB4, KCNQ2, KCNQ3, KCTD10, KLK8, LGI1, LHX6, LPPR4, MBP, MC2R, MCOLN3, MECP2, MSX2, MYO5A, NAPB, NDUFS4, NETO1, NEUROD1, NEUROD2, NID1, NOS1, NOS2, NOS3, NPAS4, NPY, NPY2R, NR4A3, NRP2, OPRD1, OTC, OTX1, PAH, PCMT1, PDE8A, PDGFRA, PDYN, PITPNA, PLAUR, PLCB1, PLCL1, PLP1, PMP22, PPT1, PRICKLE1, PRICKLE2, PRKAB1, PSAP, PTGS2, PTPRO, PURA, QKI, RAI1, RND3, RYR2, S1PR2, SCG5, SCN1A, SCN1B, SCN2B, SCN5A, SCN9A, SERPINE2, SLC10A4, SLC12A5, SLC12A6, SLC13A1, SLC17A5, SLC17A8, SLC18A3, SLC1A2, SLC1A3, SLC25A12, SLC2A1, SLC4A3, SLC7A10, SLC9A1, SLC9A6, SNAP25, SOD2, SOX1, SOX2, SPTBN2, STAM, STX1B, SUMF1, SV2A, SV2B, SYN1, SYNGAP1, SYNJ1, SZT2, TAL2, TCF12, TECTA, TEF, THRB, TPP1, TRIM2, TYRO3, UBE3A, UNC13B, USF1, USF2, USP18, VCP, ZNF24 |

| | |
|---|---|
| ADME genes | HGNC, TMEM17, SUMF1, PDE1C, CSMD1, TRIM9, C8orf34, SP140, PBX3, GRIN3A, ABCD3, SULT1B1, PON2, PDE3B, CHST3, NNMT, CYP2R1, SULF1, PON3, HAGH, SULT4A1, SLC7A8, PGRMC2, SULT1E1, TRPV4, CYP2J2, DHRS7, CYP2A6, NR1H3, DHRS1, ABCF3, ABCC4, DHRS7B, CYP7B1, ABCG8, ABCG1, CYP2D6, SLC22A2, ABCB4, ADH7, UROC1, SLC22A8, CYP7A1, NR2C2, NCOA3, CHST8, ABCG5, CYP2A13, NCOA1, CYP4F8, GSTA3, CYP2F1, INSIG1, CHST13, SLC22A6, TRPM6, VKORC1, ABCF2, NR5A2, FMO4, GSTA5, TAP2, XDH, SLCO2A1, AHRR, CYP4F11, SLCO1A2, CYP11A1, RNF40, ABCC11, NCOR1, CYP2C19, SLC22A16, GSTA2, SLC22A5, ALDH6A1, SLC22A3, ABCE1, PPARA, AKR1C3, ALDH3B2, FMO2, NR3C1, ADH1B, SLC22A4, ABCB5, CYP4F3, CYP3A7, SLC7A7, NR2C1, SLC47A1, ADH1A, AOX1, UGT8, CYP39A1, HIF1A, SQSTM1, HNF4A, ABCA5, SOD2, EAF2, ABCG2, CYP2C18, NR3C2, SLC22A15, ABCA6, GSTA1, SLC15A2, UGT2A1, NAT1, SIRT1, ABCG4, TYMS, NCOR2, SULT2A1, TAP1, ALDH2, POR, CYP46A1, IAPP, ABCA8, CABIN1, SULT2B1, TRPC1, ABCC10, CBR1, SLCO5A1, ABCD4, ALDH5A1, ESRRG, DPYD, CHST5, MGST1, GSTP1, ADH1C, NR1I3, PPARD, SLC29A2, CYP24A1, SLCO1B1, PPARGC1A, CYP2C9, FMO3, SLCO3A1, ABCC9, SLC22A7, CYP27B1, STAT3, PIAS2, SLC15A1, ADHFE1, TPMT, ALDH1A2, UGT2B10, ALDH1B1, GPX5, ALDH8A1, UGT2B4, CYP1B1, CFTR, SLCO1C1, ABCC2, MPO, CYP4F12, CAV1, ADH6, GSS, TRPC4, CYP3A5, SLC6A6, CHST12, ALDH1A1, NR1H4, FMO1, TRPM7, EPHX1, ABCA4, CHST1, SLCO2B1, SULT1A2, SLC22A9, ABCB1, GPX7, CYP3A43, SLCO1B3, CHST4, CYP4A11, ALDH1A3, ATP7B, SULT1C2, PLG, PPARG, SLC28A3, SLC22A1, TRPC6, MAT1A, CYP19A1, SLC16A1, CYP20A1, SLC47A2, SLC28A1, ABCB10, DHRS9, DDO, COMT, MGMT, CYP51A1, CBR3, UGT1A8, ALDH3A2, STK19, SLC16A7, NCOA6, ARNT, GSTM4, SLC5A12, ABCB11, CYP4F2, FOXA3, CAT, MGST2, CYP27A1, CHST11, CYP2E1, GSTO2, CYP2B6, ABCF1, CYP26C1, CREBBP, INSIG2, MGST3, GSTM3, ALDH3A1, ABCA1, UGT2B7, CYP4Z1, PDE3A, ESRRB, ABCC8, ABCC5, HSD17B11, SLC22A10, ALDH9A1, AHR, SLCO6A1, ESR2, NAT2, EPS8L3, UGT1A9, SLCO4C1, UGT1A1, UGT1A10, CYP4B1, PON1, SOD1, GPX6, CYP11B2, ESR1, TRPM1, EP300, GSR, GSTCD, SLC13A3, AKR1C2, XRCC5, SLC10A2, ABCD2, NCOA2, ABCC6, DHRS13, ABCC3, ADH5, SLC10A1, CYP11B1, UGT1A6, SLC5A6, IL6ST, NOS1, CYP17A1, DHRS7C, NR1I2, HSD11B1, SLC22A25, DHRS3, ALDH7A1, METAP1, GSTO1, EPHX2, GPX3, UGT1A3, UGT1A4, UGT1A5, CYP8B1, CHST9, UGT1A7, VDR, NR0B2, HNMT, TRPC3, ADH4, CDA, FMO5, SLC13A1, AKR1C1, CRP, CHURC1, ABCC1, CHST10, CYP2C8, GPX1, TRPC7, GSTA4, GPX2, DHRS12, SLC28A2, CYP1A2, CHST6 |

### D. Hyperparameter Settings for ML models

Table 2: Hyperparameter settings for different models used in this study.

| Model | C | Learning Rate | Trees/Iterations | Max Features | Min Sample Split | Task Penalty | Group Penalty | Max Depth |
|---|---|---|---|---|---|---|---|---|
| Log. Regression | [0.5, 0.75, 1.0, 1.25, 1.5] | - | - | - | - | - | - | - |
| SVM | [0.01, 0.1, 1, 10, 100] | - | - | - | - | - | - | - |
| Random Forest | - | - | [25, 50, 100, 125, 150] | ['sqrt', 'log2', None] | 2 | - | - | Default |
| XGBoost | - | [0.05, 0.075, 0.1, 0.15, 0.2] | [50, 75, 100, 125, 150] | - | Default | - | - | Default |
| MT-GBT | - | 0.1 | 20 | - | - | [0, 0.001, 0.01, 0.1, 0.5] | [0, 0.001, 0.01, 0.1, 0.5] | 5 |

### E. MCC values for GBT

Table 3: **MCC values with SD for Gradient-Boosted Trees.** Gradient-boosted trees were trained on all drugs and drug-specific datasets with different types of genetic information added to the dataset. The MCC values are reported with their SD on the test sets of a 5-fold cross-validation.

| Metric | All | VPA | LEV | LTG | CBZ | TPM |
|---|---|---|---|---|---|---|
| Clinical | $0.26 \pm 0.13$ | $0.22 \pm 0.23$ | $-0.09 \pm 0.13$ | $0.33 \pm 0.14$ | $0.00 \pm 0.37$ | $0.02 \pm 0.3$ |
| Clinical + Raw SNPs | $0.26 \pm 0.13$ | $0.06 \pm 0.12$ | $0.26 \pm 0.35$ | $-0.10 \pm 0.07$ | $-0.02 \pm 0.61$ | $0.08 \pm 0.23$ |
| Clinical + Gene counts | $0.24 \pm 0.05$ | $0.32 \pm 0.13$ | $-0.06 \pm 0.23$ | $-0.04 \pm 0.21$ | $0.52 \pm 0.45$ | $0.23 \pm 0.29$ |
| Clinical + Gene sets | $0.28 \pm 0.13$ | $0.19 \pm 0.1$ | $-0.02 \pm 0.21$ | $0.08 \pm 0.23$ | $-0.07 \pm 0.13$ | $-0.21 \pm 0.13$ |

### F. Upset Plots for MT Models

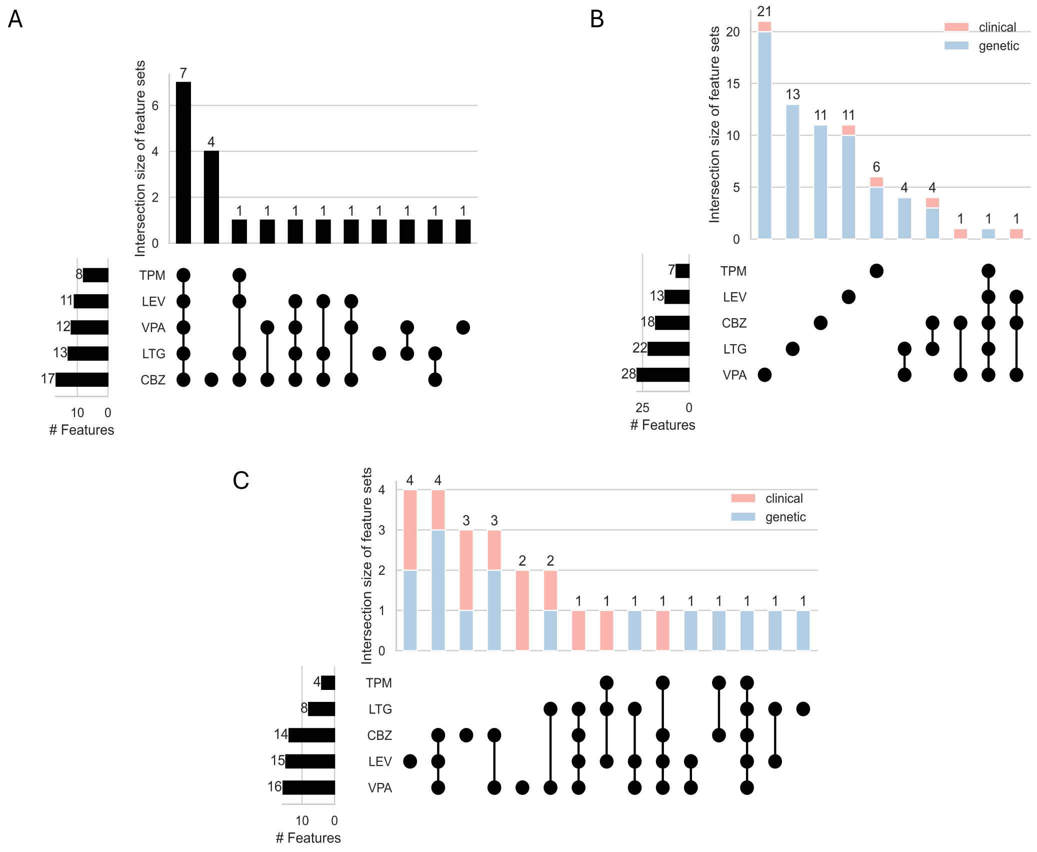


Figure 1: **Intersection of task-specific feature sets of MT gradient-boosted trees for clinical, gene counts and gene set datasets.** The feature intersections of the task-specific feature sets of the MT gradient-boosted tree model are plotted for A) the clinical dataset, B) the gene counts dataset and C) the gene set dataset. The feature set size reports the size of the task-specific feature sets and the feature intersection size shows the intersection between different combinations of task-specific feature sets. In B) and C) the colors give information about the number of clinical and genetic features shared in an intersection. The $\mu_t$ and $\mu_g$ parameter settings for the reported feature intersections are 0.5 and 0.01, respectively.
